# WOMBAT-P: Benchmarking Label-Free Proteomics Data Analysis Workflows

**DOI:** 10.1101/2023.10.02.560412

**Authors:** David Bouyssié, Pınar Altıner, Salvador Capella-Gutierrez, José M. Fernández, Yanick Paco Hagemeijer, Peter Horvatovich, Martin Hubálek, Fredrik Levander, Pierluigi Mauri, Magnus Palmblad, Wolfgang Raffelsberger, Laura Rodríguez-Navas, Dario Di Silvestre, Balázs Tibor Kunkli, Julian Uszkoreit, Yves Vandenbrouck, Juan Antonio Vizcaíno, Dirk Winkelhardt, Veit Schwämmle

## Abstract

Proteomics research encompasses a wide array of experimental designs, resulting in diverse datasets varying in structure and properties. This diversity has led to a considerable variety of software solutions for data analysis, each of them using multiple tools with different algorithms for operations like peptide-spectrum matching, protein inference, quantification, statistical analysis, and visualization. Computational workflows combine these algorithms to facilitate end-to-end analysis, spanning from raw data to detecting differentially regulated proteins. We introduce WOMBAT-P, a versatile platform designed for the automatic benchmarking and comparison of bottom-up label-free proteomics workflows. By standardizing software parameterization and workflow outputs, WOMBAT-P empowers an objective comparison of four commonly utilized data analysis workflows. Furthermore, WOMBAT-P streamlines the processing of public data based on the provided metadata, with an optional specification of 30 parameters. Wombat-P can use Sample and Data Relationship Format for Proteomics (SDRF-Proteomics) as the file input to simply process annotated local or ProteomeXchange deposited datasets. This feature offers a shortcut for data analysis and facilitates comparisons among diverse outputs. Through an examination of experimental ground truth data and a realistic biological dataset, we unveil significant disparities and a low overlap between identified and quantified proteins. WOMBAT-P not only enables rapid execution and seamless comparison of four workflows (on the same dataset) using a wide range of benchmarking metrics but also provides insights into the capabilities of different software solutions. These metrics support researchers in selecting the most suitable workflow for their specific dataset. The modular architecture of WOMBAT-P promotes extensibility and customization, making it an ideal platform for testing newly developed software tools within a realistic data analysis context.

## Introduction

Computational workflows play a crucial role in data-intensive sciences such as mass spectrometry(MS)-based proteomics by providing a way to automate and streamline complex analysis processes, and to make them easier to repeat and share with other researchers^1^. The search for an optimal data analysis solution is mostly data-dependent and cumbersome in many ways. In an optimal scenario, one would require extensive tests using ensembles of differently designed and/or parametrized workflows, all of them providing results in a comparable and standardized manner. We are aware that multiple search engines provide different results for the same dataset, and how to integrate these results is an open question. In fact, on one hand, the combination of different workflows results in a boosted false identification rate of peptides and proteins, on the other hand their intersection decreases the identification power. However, combining results from different tools remains a simple strategy to significantly improve the performance and reliability of the identification results for shotgun MS, maximizing the exploitation of the experimental MS spectra^2^.

Despite the significant developments made in recent years, there are still many challenges that must be addressed to enable more efficient, accurate, reproducible and standardized analysis of proteomic data. This holds particularly for defining, building, executing and benchmarking workflows. These challenges are not unique to proteomics, but exacerbated when compared to many other fields, due to the diversity of experimental designs, operations on data and file formats.

One major challenge is defining and identifying powerful algorithms and tools for proteomic data analysis. To achieve this, researchers require extensive knowledge of the current state-of-the-art, and software registries that provide an updated overview of the available tools such as bio.tools^3^. Additionally, benchmarks of software usage and popularity can help researchers to identify the most appropriate tools for their specific research questions^4^.

Another challenge is constructing workflows from scratch or adding new software tools to an established pipeline. This can become problematic due to file format incompatibilities, inconsistent annotation, parameter definitions, and demands on the computational environment. However, an increasing amount of format converters and shims is starting to alleviate this issue. Moreover, accurate annotation of software tool input and output, e.g. via the EDAM ontology^5^, helps identifying suitable software combinations^6,7^. Interoperability issues can be solved by using standardized file formats.

Running workflows relies on successful installations and execution settings to adapt to different computational environments. Workflow systems such as Galaxy, SnakeMake, CWL (Common Workflow Language), and Nextflow can help address these issues by being able to adapt the execution protocol to a wide range of local and cloud environments. These utilize software containers such as Docker and Singularity^8,9^ and standardized package management systems like Conda^10^ . However, many proteomics tools are still not available via fully functional software containers or as Conda packages.

Analyzing data of different origins can also pose a challenge, as it requires the raw spectra and details about the study design and experimental protocol. Standardization of this information has been initiated recently via the SDRF-Proteomics format linking data files to samples and attributes from the data acquisition^11,12^. However, this format still lacks both in details about the data analysis protocol and availability in ProteomeXchange public repositories such as the PRIDE database^13^. Moreover, very few workflows are able to directly process this standardized information.

Workflow outputs like reports on PSMs (peptide spectrum matches), peptides, proteins and differential expression/regulation can come in a myriad of different formats and depend on different levels of interpretation, making a direct and objective comparison a major challenge. Efforts to compare the output of different data analysis pipelines have been carried out quite extensively (e.g. ^14,15,16^), but they are restricted to very few datasets. Nonetheless, these studies indicated large differences between performance of various workflows, which shows the importance to benchmark different workflows and different workflow modules. Furthermore, few platforms provide benchmarking results over both different workflows and different datasets without cumbersome adaptations to work with a particular dataset.

While some efforts have been made to create central repositories for workflows, like Workflow Hub^17^ and nf-core^18^, there are still relatively few workflows listed for proteomics there. This might be due to the relatively small size of the proteomics community, as well as due to the heterogeneity and multitude of vendor specific tools and workflows often used to characterize proteomic profiles.

To address these challenges, the WOMBAT-P platform has been developed to provide a comprehensive solution for defining, executing, and comparing ensembles of workflows. The platform captures both generic and user-defined specific benchmarks of performance, efficiency, and maintainability. These benchmarks are essential for evaluating workflow performance, its components, and hardware execution. They often use reference standard datasets and evaluate performance of quantitative pre-processing and statistical analysis such as a list of differential expressed proteins, against which performance of other workflows can be measured. This ensures operation at an acceptable level and allows exploring the applicability of new software tools and their parameters over different datasets.

We demonstrate the WOMBAT-P platform with an ensemble of semantically equivalent and complete label-free quantification analysis workflows for the analysis of bottom-up proteomics data. For this purpose, we used combinations of tools known from recent literature to have a high degree of compatibility:

1. Compomics tools^19^ + FlashLFQ^20^ + MSqRob^21^ (Compomics workflow).

2. MaxQuant^22^ + NormalyzerDE^23^ (MaxQuant workflow).

3. SearchGUI^24^ + Proline^25^ + PolySTest^26^ (Proline workflow).

4. Tools from the Trans-Proteomic Pipeline (TPP)^27^ + ROTS^28^ (TPP workflow).

## Methods

### Workflow implementation

WOMBAT-P bundles different workflows for the analysis of label-free proteomics data (Fig. 1). It is built using Nextflow, a workflow language that allows running tasks across multiple computing infrastructures in a portable manner. We used an nf-core^18^ template from 2021 to set up the main framework, and we used Docker, Singularity and Apptainer containers to make installation and results maximally reproducible. The Nextflow DSL2 implementation of the platform uses one container per process, and allows describing each process in separate files, which makes it easier to maintain and update software dependencies. It also organizes the workflow steps as modules, facilitating their substitution or alternative tool combinations. As part of this study, we implemented four complete and mostly disjoint workflows based on existing tools. An overview of all processes is available in Suppl. Fig. 1. We used the release version 0.9.2.

**Figure 1:**
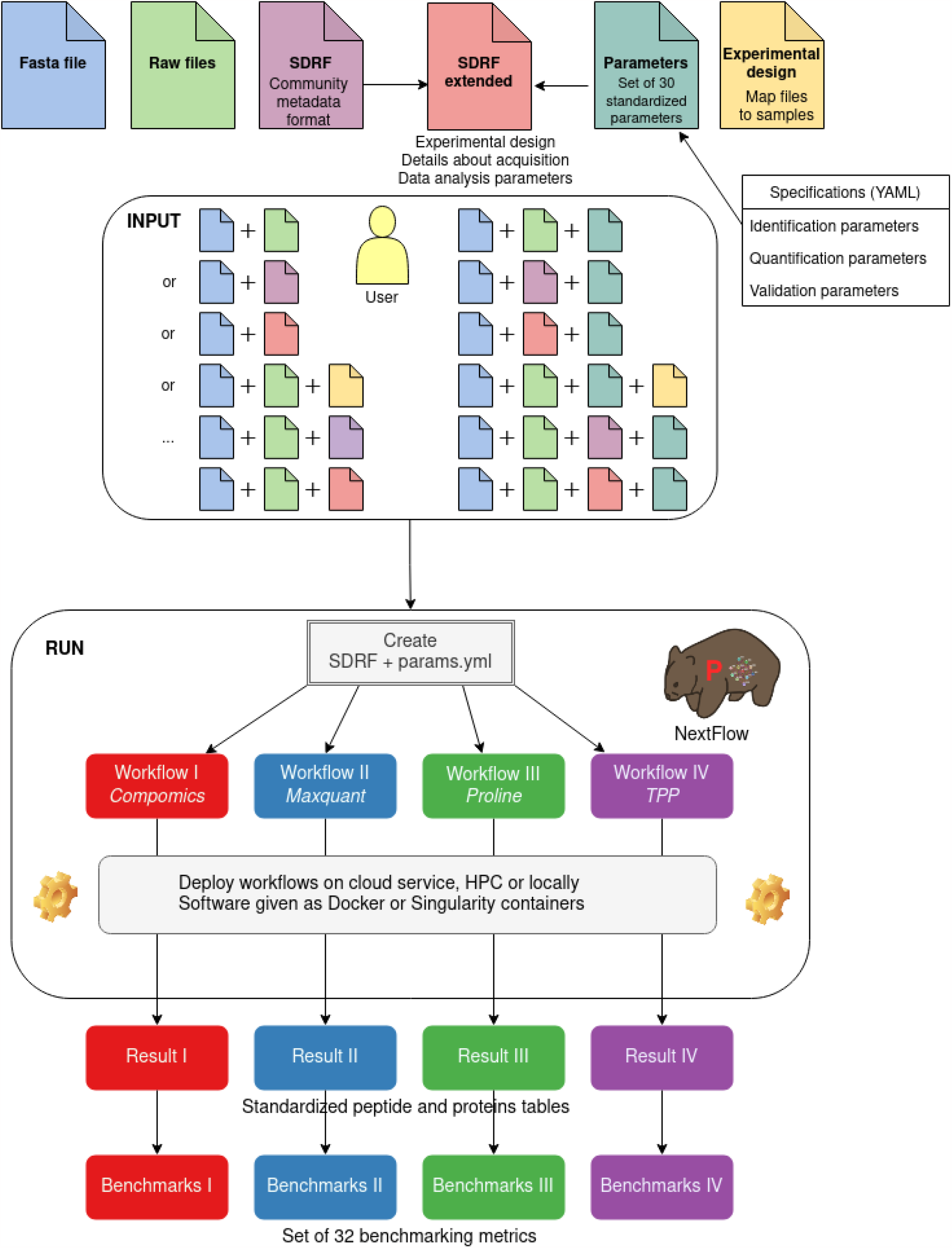
Scheme of the WOMBAT-P analysis workflows. They allow different types of input files for setting the workflow parameters and the experimental design. This can be given either in the SDRF-Proteomics file format or as separate parameter files, which can also be used to overwrite the original settings.

The first workflow (Compomics workflow) combines Compomics tools FlashLFQ^20^ and MSqRob^21^. The second workflow (MaxQuant workflow) is based on MaxQuant^22^ and NormalyzerDE^23^. In the third workflow (Proline workflow), we used SearchGUI^24^, Proline^25^, and PolySTest^26^. Finally, we also included a workflow using tools from the TPP^27^, specifically PeptideProphet^29^, ProteinProphet and StPeter^30^ with Comet^31^ for database search and ROTS^28^ for statistical analysis (TPP workflow). Multiple modules were written in Python and R scripts to facilitate conversion and parametrization within the workflows. Therein, we used MSnbase^32^ for normalization in the Compomics workflow and conversion scripts from https://github.com/jeffsocal/proteomic-id-tools in the TPP workflow.

Furthermore, we used wrProteo^33^ for in-depth analysis of the investigated ground truth dataset.

### Input options

WOMBAT-P allows a variety of input files, either based on SDRF-Proteomics annotation or via parameters given by a YAML file with the specifications of https://github.com/bigbio/proteomics-sample-metadata/blob/master/sdrf-proteomics/Data-analysis-metadata.adoc and the experimental design as tab-delimited file (see Fig. 1 for an overview of input options).

Workflow inputs are harmonized using a general set of parameters. Initialization and parameterization of the workflows are based on tools from the SDRF-pipelines^11^ and the ThermoRawFileParser^34^ for file conversion. We extended the definition of the SDRF-Proteomics data format to include a generalized set of 30 data analysis parameters (see Supplementary Table 1 and the specification of the YAML file), thus enabling the reproducibility of the data analysis via annotations with controlled vocabularies.

### Output options and benchmarks

Intermediate and final files are stored in the results folder or a folder specified via the outdir parameter. In addition to the workflow-specific output, a standardized tabular format is provided at the peptide and protein level (*stand_pep_quant_merged.csv and stand_prot_quant_merged.csv*, respectively).

For each of the workflows, WOMBAT-P calculates the same set of 32 benchmarking metrics for comparison between workflows and between different values of the data analysis parameters (see Supplementary Table 2).

Scripts for post processing are available https://github.com/wombat-p/Utilities. We used the heatmap.2 function from the gplots R package for the hierarchical clustering. For the Reactome pathway enrichment analysis of the differentially regulated proteins from each workflow^35^, we used the clusterProfiler R package^36^.

### Benchmarking Datasets

Raw data files were downloaded automatically from PRIDE by WOMBAT-P when using SDRF-Proteomics files, as the links to the files in the online repository are provided in these annotations. Fasta files for the database search were retrieved from UniProt (UniProtKB/SwissProt version Feb 3, 2023) and Sigma-Aldrich (https://www.sigmaaldrich.com/deepweb/assets/sigmaaldrich/marketing/global/fasta-files/ups1-ups2-sequences.fasta).

The ground truth dataset (PRIDE accession number PXD009815)^25^ contains a yeast background and 48 human proteins (Universal Proteomics Data Set, UPS) spiked at 10 different concentrations of UPS proteins (10 amol, 50 amol, 100 amol, 250 amol, 500 amol, fmol, 5 fmol, 10 fmol, 25 fmol and 50 fmol).

For experimental data from a standard LC-MS experiment, we used data from a study comparing COVID-19 negative and positive samples (PRIDE accession number PXD020394, from now on called COVID-19 dataset)^37^.

### Workflow registration

WOMBAT-P data provenance that can be downloaded from the WorkflowHub as an RO-Crate (Research Object Crate^38^), see https://workflowhub.eu/workflows/444. The WOMBAT-P metadata RO-Crate contains information about the workflow as well as its context. We used it to organize and share our workflow with other researchers in a standardized, interoperable, and reusable approach. The data provenance of WOMBAT-P was generated with WfExS-backend (https://github.com/inab/WfExS-backend), which is a high-level orchestrator to run scientific workflows reproducibly. It automates creating an RO-Crate by analyzing the structure and content of the computational workflow files.

Creating an RO-Crate of the WOMBAT-P workflow starts with the WOMBAT-P GitHub repository link. WfExS-backend analyzes the workflow repository, finds the workflow files and extracts the metadata, including file names, formats, file sizes, script dependencies and execution environments needed to run the workflow. The extracted metadata is mapped to the corresponding RO-Crate metadata fields following the RO-Crate specification 1.1 (https://www.researchobject.org/ro-crate/1.1) to ensure the metadata is correctly organized and represented. Then, a directory is generated using the extracted and mapped metadata with the necessary JSON-LD metadata files, including the ro-crate-metadata.json file, which contains the comprehensive metadata for the RO-Crate. In addition, the directory includes the workflow files and associated data to ensure all relevant files are included and can be accessed.

## Availability

All workflows and documentation are available in GitHub at https://github.com/wombat-p/WOMBAT-Pipelines under the MIT license.

The main result files and the files needed for running the workflows are deposited at https://github.com/wombat-p/WOMBAT-P_Processed.

## Results

We provide detailed evaluations of the workflows and their results on the basis of two different datasets. WOMBAT version 0.9.2 was used for analyzing these datasets. The first dataset is a ground truth dataset with known information about the expected quantitative changes, and thus serves to directly assess workflow performance. The second dataset comes from a typical label-free proteomics experiment and thus should resemble the structure and properties of such.

### Ground truth dataset

Comparison of different data analysis software and pipelines was performed on data from a ground-truth spike-in experiment^25^.

Regarding the overall number of identified peptides and proteins, we observed that TPP reported a much lower peptide count than *MaxQuant* and Compomics (Fig. 2A). We suspect that this is due to the prior filtering of peptides for protein FDRs (False Discovery Rate) in the TPP workflow. This also shows that the comparison at the same stage in the analysis is often hampered by even slightly different data treatment in the workflows. Despite the differences in peptide quantifications, all workflows showed similar numbers of proteins (Fig. 2A).

**Figure 2:**
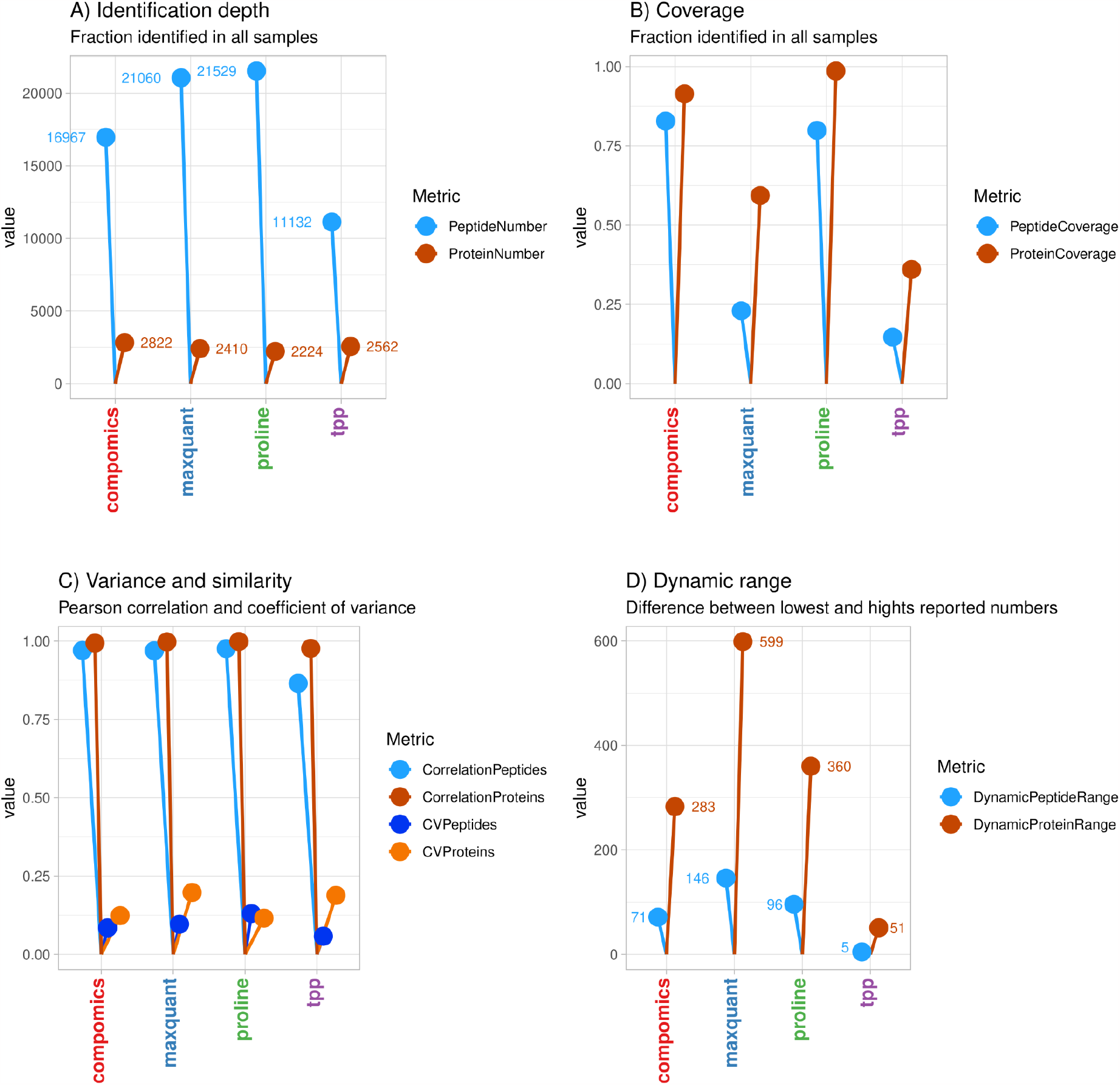
Summary of the benchmarking results for the ground truth dataset.

### Proline and FlashLFQ from the Compomics

workflow reported higher coverage of peptides and proteins across samples compared to TPP, which lacked a match-between-runs option (Fig 2B). A lower coverage of quantified peptides across samples was observed in MaxQuant, and even lower in TPP. The lower number in TPP is likely due to the inability to use information from other MS runs to improve overall peptide and protein identifications (Fig. 2B).

Higher coverage decreased the variation of protein abundance values between replicates, and we consistently observed high correlations for all workflows, with the exception of TPP, which provided slightly lower performance (Fig. 2C).

Finally, we evaluated the dynamic range, i.e. the range of reported quantitative values for identified peptides and proteins by the workflows, to reproduce actual changes in protein and peptide abundance within the mass spectrometer’s sensitivity. We found that *MaxQuant* had an about 10-fold higher range of protein abundance changes of 599 when compared to TPP with 51, which is considerably different from the variability observed in the other workflows (Fig. 2D). Here, StPeter from TPP uses a different method of quantification than the other tools. It is noteworthy to mention that a higher dynamic range does not necessarily reflect a more accurate output as distinct summarization of PSMs and peptides can lead to different deviations from the linear response.

While generic benchmarking metrics such as the number of peptide and protein identifications offer valuable insights into workflow performance, it is equally important to assess the overlap and similarity between the obtained results. We then evaluated the overlap in protein identifications across all samples and workflows (Suppl. Fig. 2), revealing distinct patterns both across and within the four workflows. The Compomics and Proline workflows exhibited a higher overlap within different samples.

In terms of quantified peptides, we observed that their similarity with respect to their relative abundances was generally higher within each workflow (Suppl. Fig. 3), likely due to variations in the methods used to quantify peptide and protein intensities. This trend was particularly noticeable in the TPP workflow, where quantification was based on StPeter, an algorithm that incorporates spectral counting. Furthermore, the workflows successfully grouped samples with the same UPS protein concentrations in most instances.

The samples of the experiment consist of 48 human proteins spiked at levels of in yeast proteome. Thus, only all proteins annotated as this species (human in this dataset) are expected to vary between samples, while proteins of the other species (yeast in this dataset) are expected to be always detected at a constant level.

We run the workflows with median normalization, i.e. without using a specialized normalization method that adapt to the rather particular experimental design. Given the large concentration changes of the spike-in UPS proteins, this design led to an apparent under-expression of the background proteins in the conditions where the UPS proteins are highly abundant, thus providing misleading quantitative changes of the background proteins. A specialized analysis and extensive interpretations for this spike-in ground truth data including use of appropriate normalization is available in the supplemental material using wrProteo.

However, in a regular experiment it is not known in advance which proteins are expected to be constant in abundance. Therefore, we did not apply such data transformations in order to provide an objective view on workflow performance. This also meant that the “official” ground truth of finding 48 differentially regulated UPS proteins became mixed with differentially regulated yeast proteins. While assuming that this should not affect the outcome too drastically, we find surprisingly different results when comparing the output from the four WOMBAT-P workflows.

We checked how well the workflows detected the UPS proteins as differentially regulated in a challenging case of their low concentrations being 10 amol versus 500 amol. For that, we assessed the number of human proteins observed as variants (providing the sensitivity) and the percentage of wrongly detected yeast variants (providing the specificity). When examining the outputs of the workflows in terms of differentially regulated proteins (FDR < 0.05), there was a poor agreement among all workflows (Fig. 3). The Proline and Compomics workflows showed the highest number of correctly identified UPS proteins with a total of 37 and 35 proteins, respectively. Notably, 14 of the 37 UPS proteins identified by Proline as significantly different were only found by this particular workflow. Similarly, the Compomics workflow reported 12 UPS proteins uniquely found for this workflow. When comparing with the number of differentially regulated yeast proteins representing false positives, their numbers were lower and there was no overlap between workflows. Notably, all eight proteins found in at least two workflows were UPS proteins. For a more systematic exploration of the ground truth, we refer to Supplementary File 1.

**Figure 3:**
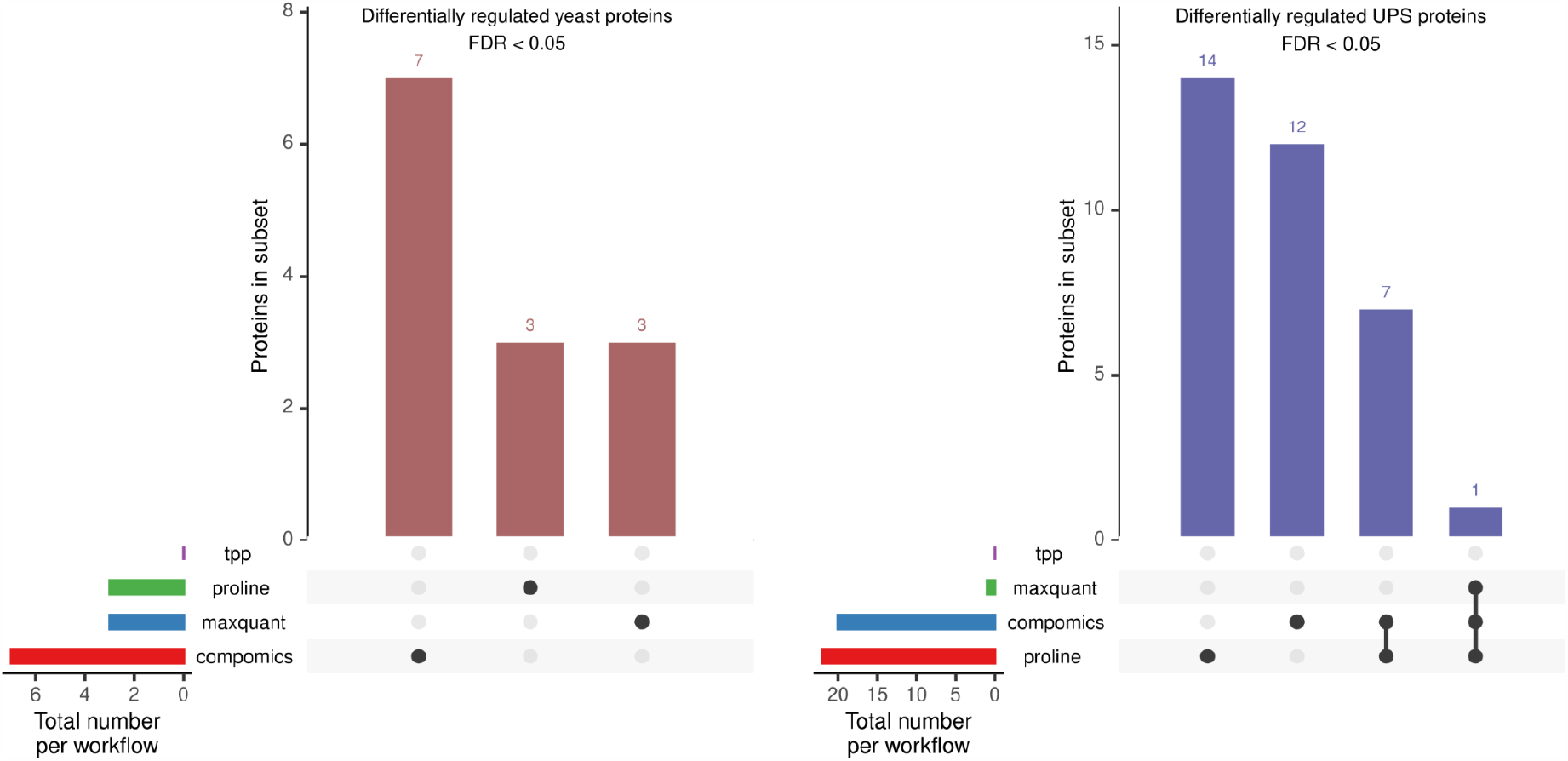
Comparison of proteins found to be differentially regulated between samples with UPS amounts of 10 amol and 500 amol, and a background of equally abundant yeast proteins. Differential regulation was determined by the respective workflow components for statistical testing, which uses different statistical approaches.

We also measured the usage of available computational resources with different metrics (Figure 4). These metrics were extracted from a trace report which was computed by nextflow. It contains information about each executed process in the pipeline. The results varied widely across different tasks, which could be attributed to different implementations and differently assigned subtasks of the major data transformations.

**Figure 4:**
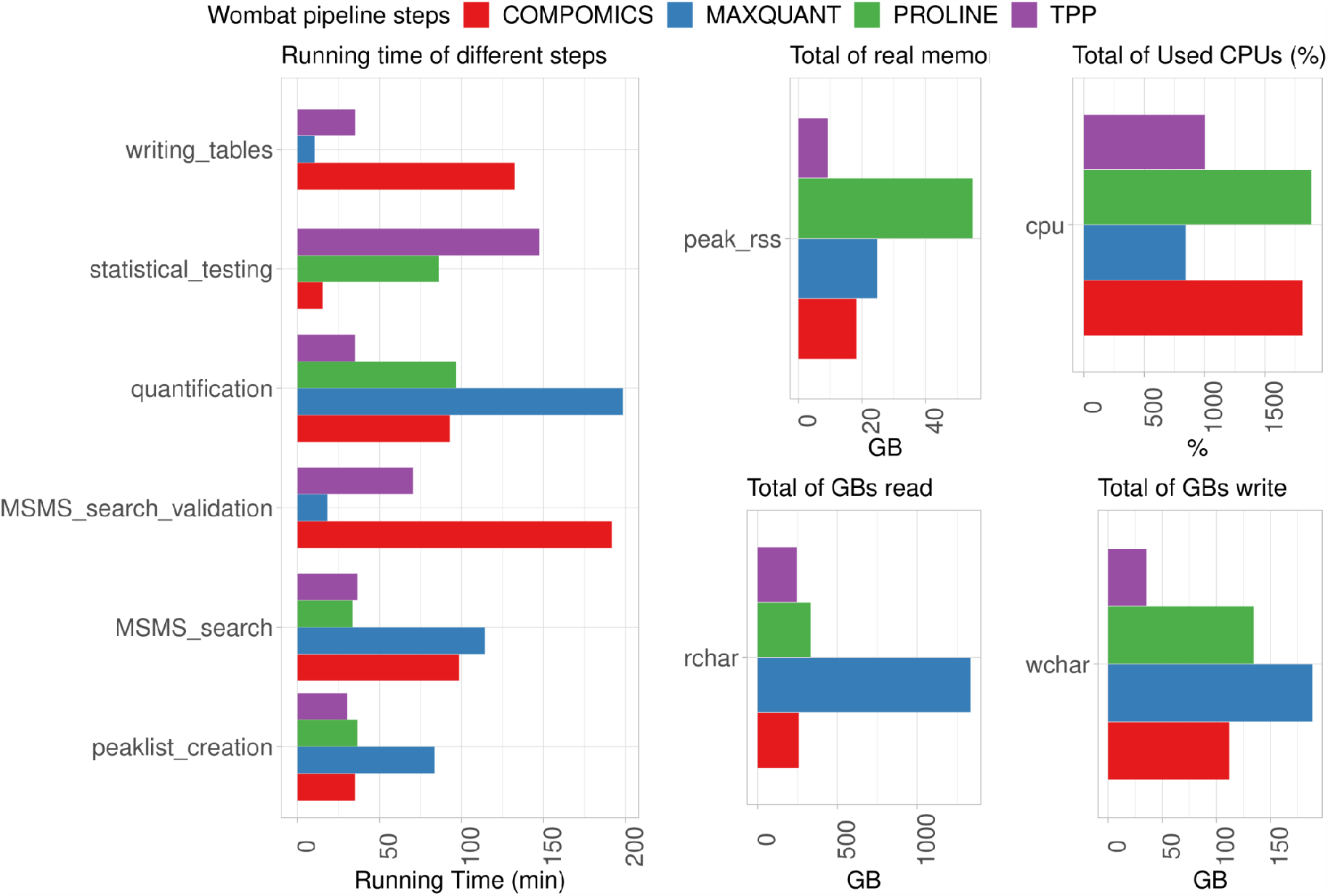
Comparison of used computational resources for the different workflows. Given that *MaxQuant* bundles multiple operations, we only report running times which were available in the *MaxQuant* output. rchar: amount of the data being read, wchar: amount of written data, peak_rss: peak amount of RAM, cpu: average number of used CPU threads in percentage.

On the whole, we noticed that *Proline* and *Compomics* used CPUs more efficiently and required more memory, while the *MaxQuant* workflow performed more file read/write operations. In summary, when utilizing the ground truth dataset with UPS proteins spiked into a yeast background, significant differences were observed among the outputs of the various workflows.

### Performance on biological data from COVID-19 study

Ground truth datasets can be limited in accurately replicating biological samples. Therefore, we decided to also assess the performance of WOMBAT-P using a recent dataset that compared COVID-19 positive and negative subjects^37^. The utilization of this dataset led to slightly different results, showing that workflow performance can depend strongly on each particular dataset it processes.

When comparing the different benchmarks (Fig. 5), the *Compomics* and TPP workflows consistently exhibited lower peptide quantification numbers, whereas the Compomics workflow demonstrated up to 400 more quantified proteins, when compared to the other workflows. When evaluating the variance and correlation levels within replicates of the same sample type, the correlations were lower compared to the ground truth data. This disparity can be attributed to the increased biological variability among the samples (from controls and patients) and to the lower number of identified proteins. Notably, the TPP workflow exhibited the lowest coefficient of variance and the highest protein abundance value correlations of non-log-transformed values. It is worth mentioning that we observed significant differences in the dynamic ranges of protein and peptide quantifications, similar to what was observed in the ground truth dataset. These differences could have influenced the high correlation values and the low coefficient of variance for the TPP workflow.

**Figure 5:**
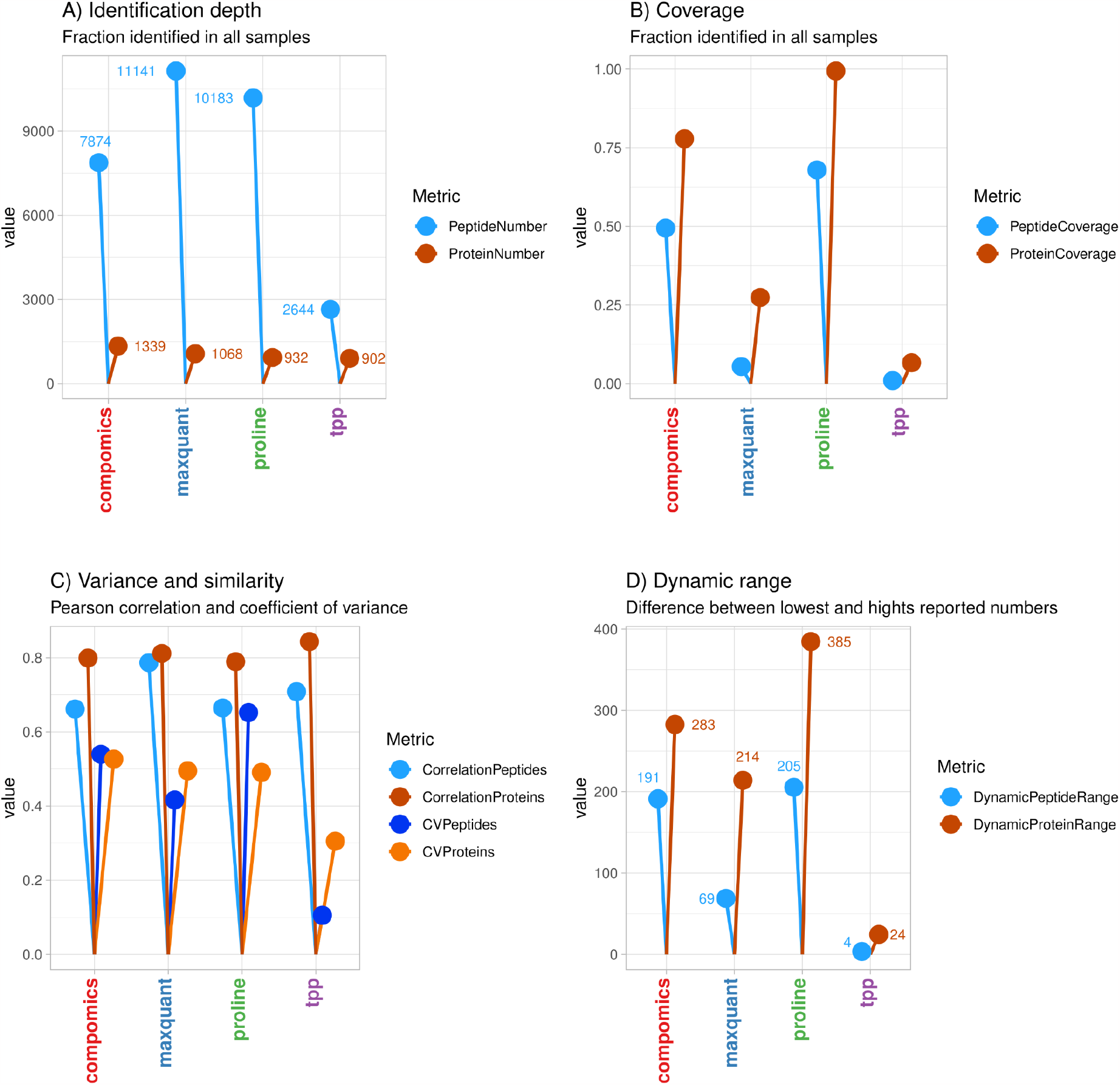
Summary of the benchmarking results for biological dataset.

Run times showed a diverse picture, similarly to what was observed for the UPS dataset (Suppl. Fig. 4). Additionally, computer resources used for the processing of this dataset exhibited very close relative values, compared to the UPS dataset.

When comparing the overlap of identified proteins within and across the workflows (Suppl. Fig. 5), it could be confirmed that there was a much higher similarity in the proteomes measured amongst samples analyzed with the same workflow. This can not be merely attributed to applied match-between run options, given that the *TPP* workflow separates similarly. While it is well attested in the literature^39,40,41^ that different search engines identify non-identical sets of peptides from the same data, downstream components, such as for PSM validation^42^ or protein inference^43^ may also be of considerable influence.

The quantitative comparison of the workflows’ results (Fig. 6) provided more insights into workflow performance. For the correlations within the samples of a given workflow, the Proline workflow performed with a highest similarity within all samples. Notably, the results from *Compomics* and *MaxQuant* workflows were sufficiently similar to separate the two sample types between them.

**Fig. 6:**
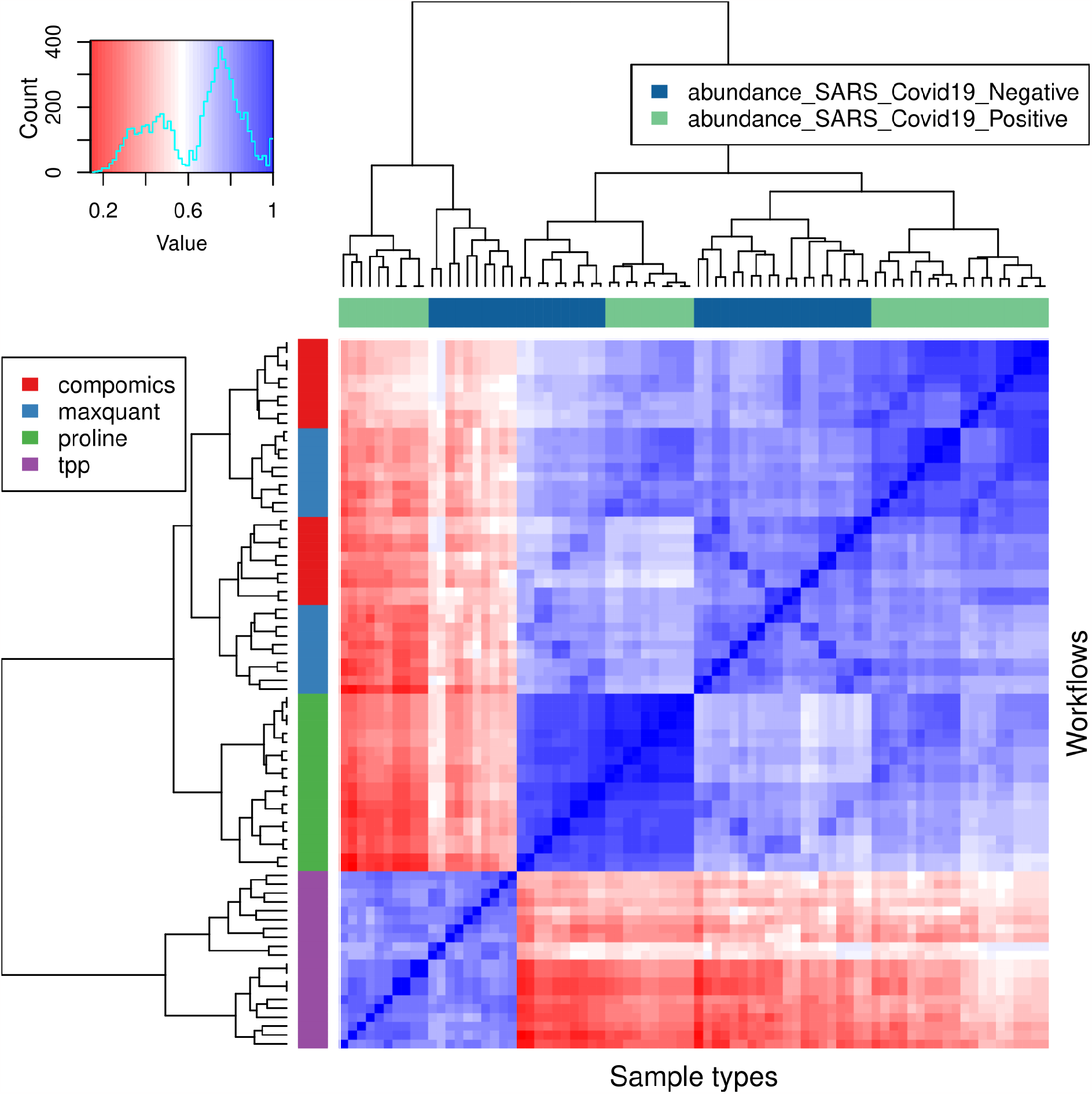
Similarity of workflow results using the COVID-19 dataset. Pearson correlation values were calculated to compare the similarity on a quantitative level, and then the correlations were arranged using hierarchical clustering. In contrast to the benchmarking metrics in Figs. and 5, here the Pearson correlation values were calculated from the log_2_-transformed values.

We furthermore compared the protein groups found to be differentially regulated (Suppl. Fig. 6). While more than one hundred proteins were detected in at least three of the four workflows, they disagreed to a high degree, leading to more than 500 proteins being uniquely found to be significantly changing between the COVID-19 positive and negative samples. However, further analysis via Reactome pathway enrichment led to considerable agreement between the results of Compomics, Maxquant and Proline workflows for the most enriched pathways (Suppl. Fig. 7).

In summary, we found both similarities and differences when comparing workflow performances. The noise levels, as expected, were higher for the COVID-19 dataset and showed higher similarity between the different workflows (Figs. 2C and 5C).

## Discussion

This study introduces WOMBAT-P as an innovative solution for addressing the challenges of large-scale proteomic data analysis. It relies on the importance of automated data quality control and validation in scaling up to analyzing a large number of files. The scalability of WOMBAT-P using high-performance computing (HPC) environments and its utilization of software containers enable reproducible analyses, making it a valuable tool for both public and in-house label-free data analysis at a large scale.

One notable feature of WOMBAT-P is its ability to create SDRF-Proteomics file templates and produce harmonized outputs, facilitating easy benchmarking and comparison of results. This comprehensive approach enhances the robustness and reliability of proteomic data analysis.

We found different performances of the workflows when testing them on two selected datasets. *TPP* showed relatively lower protein and peptide identification numbers, likely due to the absence of a match-between-runs feature in this particular workflow. However, the workflow still showed considerable performance. This finding suggests the absence of a universally optimal solution for proteomic data analysis. It underscores the significance of data-driven workflow analysis using benchmarking metrics to assess and identify the best-performing solutions for proteomic data analysis.

The modularized architecture of WOMBAT-P enables the incorporation of new processing algorithms and steps and workflows, including new advancements such as tools applying deep learning and further downstream software for biological interpretation. This flexibility enhances the capabilities of WOMBAT-P and expands its potential for further developments in proteomic analyses.

We view WOMBAT-P as a continuously evolving platform with ongoing development and open development, with new modules and updates anticipated in the near future, and therefore invite for contributions from the proteomics bioinformatics community. Achieving exchangeability of certain operations like the downstream statistical tests across all four workflows will make WOMBAT-P a more versatile and flexible tool for proteomic data analysis.

In conclusion, this study highlights the significance and relevance of WOMBAT-P for proteomic data analysis. By providing a comprehensive tool that enhances accuracy, scalability, and comparability in the analysis of large-scale proteomic analyses, WOMBAT-P addresses critical challenges and contributes to advancing our understanding of complex biological systems.

We are aware that benchmarking of software can be complex, due to the number of options available in terms of parameters, the different characteristics of the benchmarking datasets and the need to be expert in all the tools used in the benchmarking. WOMBAT-P allows users to run and compare different configurations and thus explore alternative and more optimized setups.

## Acknowledgements

This work was funded by ELIXIR (https://elixir-europe.org/), the European research infrastructure for life-science data.

## References

(1) Harjes, J.; Link, A.; Weibulat, T.; Triebel, D.; Rambold, G. FAIR Digital Objects in Environmental and Life Sciences Should Comprise Workflow Operation Design Data and Method Information for Repeatability of Study Setups and Reproducibility of Results. Database 2020, 2020. 10.1093/database/baaa059.

(2) Zhao, P.; Zhong, J.; Liu, W.; Zhao, J.; Zhang, G. Protein-Level Integration Strategy of Multiengine MS Spectra Search Results for Higher Confidence and Sequence Coverage. J. Proteome Res. 2017, 16 (12), 4446–4454.

(3) Ison, J.; Ienasescu, H.; Chmura, P.; Rydza, E.; Ménager, H.; Kalaš, M.; Schwämmle, V.; Grüning, B.; Beard, N.; Lopez, R.; Duvaud, S.; Stockinger, H.; Persson, B.; Vařeková, R. S.; Raček, T.; Vondrášek, J.; Peterson, H.; Salumets, A.; Jonassen, I.; Hooft, R.; Nyrönen, T.; Valencia, A.; Capella, S.; Gelpí, J.; Zambelli, F.; Savakis, B.; Leskošek, B.; Rapacki, K.; Blanchet, C.; Jimenez, R.; Oliveira, A.; Vriend, G.; Collin, O.; van Helden, J.; Løngreen, P.; Brunak, S. The Bio.tools Registry of Software Tools and Data Resources for the Life Sciences. Genome Biol. 2019, 20 (1), 164.

(4) Capella-Gutierrez, S.; Iglesia, D. de la; Haas, J.; Lourenco, A.; Fernández, J. M.; Repchevsky, D.; Dessimoz, C.; Schwede, T.; Notredame, C.; Gelpi, J. L.; Valencia, A. Lessons Learned: Recommendations for Establishing Critical Periodic Scientific Benchmarking. bioRxiv, 2017. 10.1101/181677.

(5) Ison, J.; Kalas, M.; Jonassen, I.; Bolser, D.; Uludag, M.; McWilliam, H.; Malone, J.; Lopez, R.; Pettifer, S.; Rice, P. EDAM: An Ontology of Bioinformatics Operations, Types of Data and Identifiers, Topics and Formats. Bioinformatics 2013, 29 (10), 1325–1332.

(6) Palmblad, M.; Lamprecht, A.-L.; Ison, J.; Schwämmle, V. Automated Workflow Composition in Mass Spectrometry-Based Proteomics. Bioinformatics 2019, 35 (4), 656–664.

(7) Kasalica, V.; Schwämmle, V.; Palmblad, M.; Ison, J.; Lamprecht, A.-L. APE in the Wild: Automated Exploration of Proteomics Workflows in the Bio.tools Registry. J. Proteome Res. 2021, 20 (4), 2157–2165.

(8) da Veiga Leprevost, F.; Grüning, B. A.; Alves Aflitos, S.; Röst, H. L.; Uszkoreit, J.; Barsnes, H.; Vaudel, M.; Moreno, P.; Gatto, L.; Weber, J.; Bai, M.; Jimenez, R. C.; Sachsenberg, T.; Pfeuffer, J.; Vera Alvarez, R.; Griss, J.; Nesvizhskii, A. I.; Perez-Riverol, Y. BioContainers: An Open-Source and Community-Driven Framework for Software Standardization. Bioinformatics 2017, 33 (16), 2580–2582.

(9) Bai, J.; Bandla, C.; Guo, J.; Vera Alvarez, R.; Bai, M.; Vizcaíno, J. A.; Moreno, P.; Grüning, B.; Sallou, O.; Perez-Riverol, Y. BioContainers Registry: Searching Bioinformatics and Proteomics Tools, Packages, and Containers. J. Proteome Res. 2021, 20 (4), 2056–2061.

(10) Grüning, B.; Dale, R.; Sjödin, A.; Chapman, B. A.; Rowe, J.; Tomkins-Tinch, C. H.; Valieris, R.; Köster, J.; Bioconda Team. Bioconda: Sustainable and Comprehensive Software Distribution for the Life Sciences. Nat. Methods 2018, 15 (7), 475–476.

(11) Perez-Riverol, Y.; European Bioinformatics Community for Mass Spectrometry. Toward a Sample Metadata Standard in Public Proteomics Repositories. J. Proteome Res. 2020, 19 (10), 3906–3909.

(12) Dai, C.; Füllgrabe, A.; Pfeuffer, J.; Solovyeva, E. M.; Deng, J.; Moreno, P.; Kamatchinathan, S.; Kundu, D. J.; George, N.; Fexova, S.; Grüning, B.; Föll, M. C.; Griss, J.; Vaudel, M.; Audain, E.; Locard-Paulet, M.; Turewicz, M.; Eisenacher, M.; Uszkoreit, J.; Van Den Bossche, T.; Schwämmle, V.; Webel, H.; Schulze, S.; Bouyssié, D.; Jayaram, S.; Duggineni, V. K.; Samaras, P.; Wilhelm, M.; Choi, M.; Wang, M.; Kohlbacher, O.; Brazma, A.; Papatheodorou, I.; Bandeira, N.; Deutsch, E. W.; Vizcaíno, J. A.; Bai, M.; Sachsenberg, T.; Levitsky, L. I.; Perez-Riverol, Y. A Proteomics Sample Metadata Representation for Multiomics Integration and Big Data Analysis. Nat. Commun. 2021, 12 (1), 5854.

(13) Perez-Riverol, Y.; Bai, J.; Bandla, C.; García-Seisdedos, D.; Hewapathirana, S.; Kamatchinathan, S.; Kundu, D. J.; Prakash, A.; Frericks-Zipper, A.; Eisenacher, M.; Walzer, M.; Wang, S.; Brazma, A.; Vizcaíno, J. A. The PRIDE Database Resources in 2022: A Hub for Mass Spectrometry-Based Proteomics Evidences. Nucleic Acids Res. 2022, 50 (D1), D543–D552.

(14) Lietzén, N.; Natri, L.; Nevalainen, O. S.; Salmi, J.; Nyman, T. A. Compid: A New Software Tool to Integrate and Compare MS/MS Based Protein Identification Results from Mascot and Paragon. J. Proteome Res. 2010, 9 (12), 6795–6800.

(15) Hoekman, B.; Breitling, R.; Suits, F.; Bischoff, R.; Horvatovich, P. msCompare: A Framework for Quantitative Analysis of Label-Free LC-MS Data for Comparative Candidate Biomarker Studies. Mol. Cell. Proteomics 2012, 11 (6), M111.015974.

(16) Locard-Paulet, M.; Bouyssié, D.; Froment, C.; Burlet-Schiltz, O.; Jensen, L. J. Comparing 22 Popular Phosphoproteomics Pipelines for Peptide Identification and Site Localization. J. Proteome Res. 2020, 19 (3), 1338–1345.

(17) Goble, C.; Soiland-Reyes, S.; Bacall, F.; Owen, S.; Williams, A.; Eguinoa, I.; Droesbeke, B.; Leo, S.; Pireddu, L.; Rodríguez-Navas, L.; Fernández, J. M.; Capella-Gutierrez, S.; Ménager, H.; Grüning, B.; Serrano-Solano, B.; Ewels, P.; Coppens, F. Implementing FAIR Digital Objects in the EOSC-Life Workflow Collaboratory. Zenodo 2021. 10.5281/ZENODO.4605654.

(18) Ewels, P. A.; Peltzer, A.; Fillinger, S.; Patel, H.; Alneberg, J.; Wilm, A.; Garcia, M. U.; Di Tommaso, P.; Nahnsen, S. The Nf-Core Framework for Community-Curated Bioinformatics Pipelines. Nat. Biotechnol. 2020, 38 (3), 276–278.

(19) Barsnes, H.; Vaudel, M.; Colaert, N.; Helsens, K.; Sickmann, A.; Berven, F. S.; Martens, L. Compomics-Utilities: An Open-Source Java Library for Computational Proteomics. BMC Bioinformatics 2011, 12, 70.

(20) Millikin, R. J.; Shortreed, M. R.; Scalf, M.; Smith, L. M. Fast, Free, and Flexible Peptide and Protein Quantification with FlashLFQ. Methods Mol. Biol. 2023, 2426, 303–313.

(21) Goeminne, L. J. E.; Gevaert, K.; Clement, L. Experimental Design and Data-Analysis in Label-Free Quantitative LC/MS Proteomics: A Tutorial with MSqRob. J. Proteomics 2018, 171, 23–36.

(22) Tyanova, S.; Temu, T.; Cox, J. The MaxQuant Computational Platform for Mass Spectrometry-Based Shotgun Proteomics. Nat. Protoc. 2016, 11 (12), 2301–2319.

(23) Willforss, J.; Chawade, A.; Levander, F. NormalyzerDE: Online Tool for Improved Normalization of Omics Expression Data and High-Sensitivity Differential Expression Analysis. J. Proteome Res. 2019, 18 (2), 732–740.

(24) Barsnes, H.; Vaudel, M. SearchGUI: A Highly Adaptable Common Interface for Proteomics Search and de Novo Engines. J. Proteome Res. 2018, 17 (7), 2552–2555.

(25) Bouyssié, D.; Hesse, A.-M.; Mouton-Barbosa, E.; Rompais, M.; Macron, C.; Carapito, C.; Gonzalez de Peredo, A.; Couté, Y.; Dupierris, V.; Burel, A.; Menetrey, J.-P.; Kalaitzakis, A.; Poisat, J.; Romdhani, A.; Burlet-Schiltz, O.; Cianférani, S.; Garin, J.; Bruley, C. Proline: An Efficient and User-Friendly Software Suite for Large-Scale Proteomics. Bioinformatics 2020, 36 (10), 3148–3155.

(26) Schwämmle, V.; Hagensen, C. E.; Rogowska-Wrzesinska, A.; Jensen, O. N. PolySTest: Robust Statistical Testing of Proteomics Data with Missing Values Improves Detection of Biologically Relevant Features. Mol. Cell. Proteomics 2020, 19 (8), 1396–1408.

(27) Deutsch, E. W.; Mendoza, L.; Shteynberg, D. D.; Hoopmann, M. R.; Sun, Z.; Eng, J. K.; Moritz, R. L. Trans-Proteomic Pipeline: Robust Mass Spectrometry-Based Proteomics Data Analysis Suite. J. Proteome Res. 2023, 22 (2), 615–624.

(28) Suomi, T.; Seyednasrollah, F.; Jaakkola, M. K.; Faux, T.; Elo, L. L. ROTS: An R Package for Reproducibility-Optimized Statistical Testing. PLoS Comput. Biol. 2017, 13 (5), e1005562.

(29) Ma, K.; Vitek, O.; Nesvizhskii, A. I. A Statistical Model-Building Perspective to Identification of MS/MS Spectra with PeptideProphet. BMC Bioinformatics 2012, 13 Suppl 16 (Suppl 16), S1.

(30) Hoopmann, M. R.; Winget, J. M.; Mendoza, L.; Moritz, R. L. StPeter: Seamless Label-Free Quantification with the Trans-Proteomic Pipeline. J. Proteome Res. 2018, 17 (3), 1314–1320.

(31) Eng, J. K.; Jahan, T. A.; Hoopmann, M. R. Comet: An Open-Source MS/MS Sequence Database Search Tool. Proteomics 2013, 13 (1), 22–24.

(32) Gatto, L.; Gibb, S.; Rainer, J. MSnbase, Efficient and Elegant R-Based Processing and Visualization of Raw Mass Spectrometry Data. J. Proteome Res. 2021, 20 (1), 1063–1069.

(33) Raffelsberger, W. wrProteo: Proteomics Data Analysis Functions; 2023.

(34) Hulstaert, N.; Shofstahl, J.; Sachsenberg, T.; Walzer, M.; Barsnes, H.; Martens, L.; Perez-Riverol, Y. ThermoRawFileParser: Modular, Scalable, and Cross-Platform RAW File Conversion. J. Proteome Res. 2020, 19 (1), 537–542.

(35) Gillespie, M.; Jassal, B.; Stephan, R.; Milacic, M.; Rothfels, K.; Senff-Ribeiro, A.; Griss, J.; Sevilla, C.; Matthews, L.; Gong, C.; Deng, C.; Varusai, T.; Ragueneau, E.; Haider, Y.; May, B.; Shamovsky, V.; Weiser, J.; Brunson, T.; Sanati, N.; Beckman, L.; Shao, X.; Fabregat, A.; Sidiropoulos, K.; Murillo, J.; Viteri, G.; Cook, J.; Shorser, S.; Bader, G.; Demir, E.; Sander, C.; Haw, R.; Wu, G.; Stein, L.; Hermjakob, H.; D’Eustachio, P. The Reactome Pathway Knowledgebase 2022. Nucleic Acids Res. 2022, 50 (D1), D687–D692.

(36) Wu, T.; Hu, E.; Xu, S.; Chen, M.; Guo, P.; Dai, Z.; Feng, T.; Zhou, L.; Tang, W.; Zhan, L.; Fu, X.; Liu, S.; Bo, X.; Yu, G. clusterProfiler 4.0: A Universal Enrichment Tool for Interpreting Omics Data. Innovation (Camb) 2021, 2 (3), 100141.

(37) Rivera, B.; Leyva, A.; Portela, M. M.; Moratorio, G.; Moreno, P.; Durán, R.; Lima, A. Quantitative Proteomic Dataset from Oro- and Naso-Pharyngeal Swabs Used for COVID-19 Diagnosis: Detection of Viral Proteins and Host’s Biological Processes Altered by the Infection. Data Brief 2020, 32, 106121.

(38) Soiland-Reyes, S.; Sefton, P.; Crosas, M.; Castro, L. J.; Coppens, F.; Fernández, J. M.; Garijo, D.; Grüning, B.; La Rosa, M.; Leo, S.; Ó Carragáin, E.; Portier, M.; Trisovic, A.; RO-Crate Community; Groth, P.; Goble, C. Packaging Research Artefacts with RO-Crate. Data Sci. 2022, 5 (2), 97–138.

(39) Mohammed, Y.; Palmblad, M. Visualizing and Comparing Results of Different Peptide Identification Methods. Brief. Bioinform. 2018, 19 (2), 210–218.

(40) De La Toba, E. A.; Anapindi, K. D. B.; Sweedler, J. V. Assessment and Comparison of Database Search Engines for Peptidomic Applications. J. Proteome Res. 2023. 10.1021/acs.jproteome.2c00307.

(41) Yuan, Z.-F.; Lin, S.; Molden, R. C.; Garcia, B. A. Evaluation of Proteomic Search Engines for the Analysis of Histone Modifications. J. Proteome Res. 2014, 13 (10), 4470–4478.

(42) Shteynberg, D.; Deutsch, E. W.; Lam, H.; Eng, J. K.; Sun, Z.; Tasman, N.; Mendoza, L.; Moritz, R. L.; Aebersold, R.; Nesvizhskii, A. I. iProphet: Multi-Level Integrative Analysis of Shotgun Proteomic Data Improves Peptide and Protein Identification Rates and Error Estimates. Mol. Cell. Proteomics 2011, 10 (12), M111.007690.

(43) Audain, E.; Uszkoreit, J.; Sachsenberg, T.; Pfeuffer, J.; Liang, X.; Hermjakob, H.; Sanchez, A.; Eisenacher, M.; Reinert, K.; Tabb, D. L.; Kohlbacher, O.; Perez-Riverol, Y. In-Depth Analysis of Protein Inference Algorithms Using Multiple Search Engines and Well-Defined Metrics. J. Proteomics 2017, 150, 170–182.

