## Supplementary Figures for "WOMBAT-P: Benchmarking Label-Free Proteomics Data Analysis Workflows"

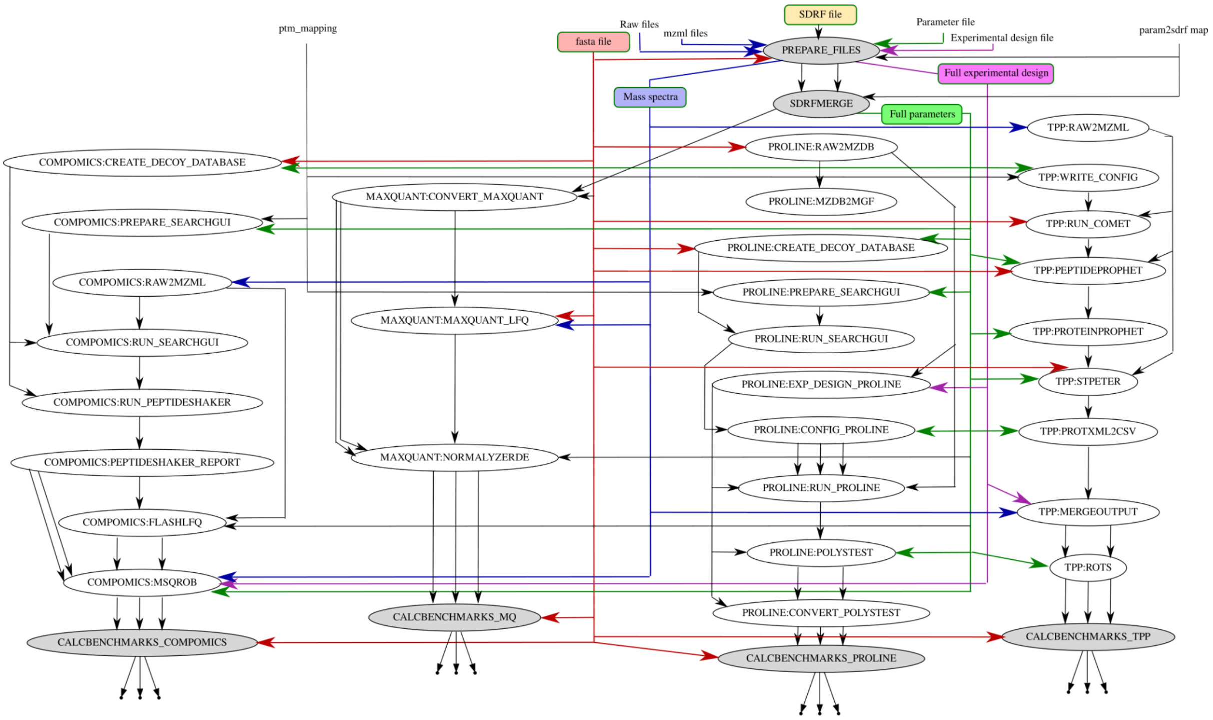


Suppl. Fig. 1: Flowchart of all processes run by WOMBAT-P. The arrows describe the data flow from input files to output files and benchmarks.

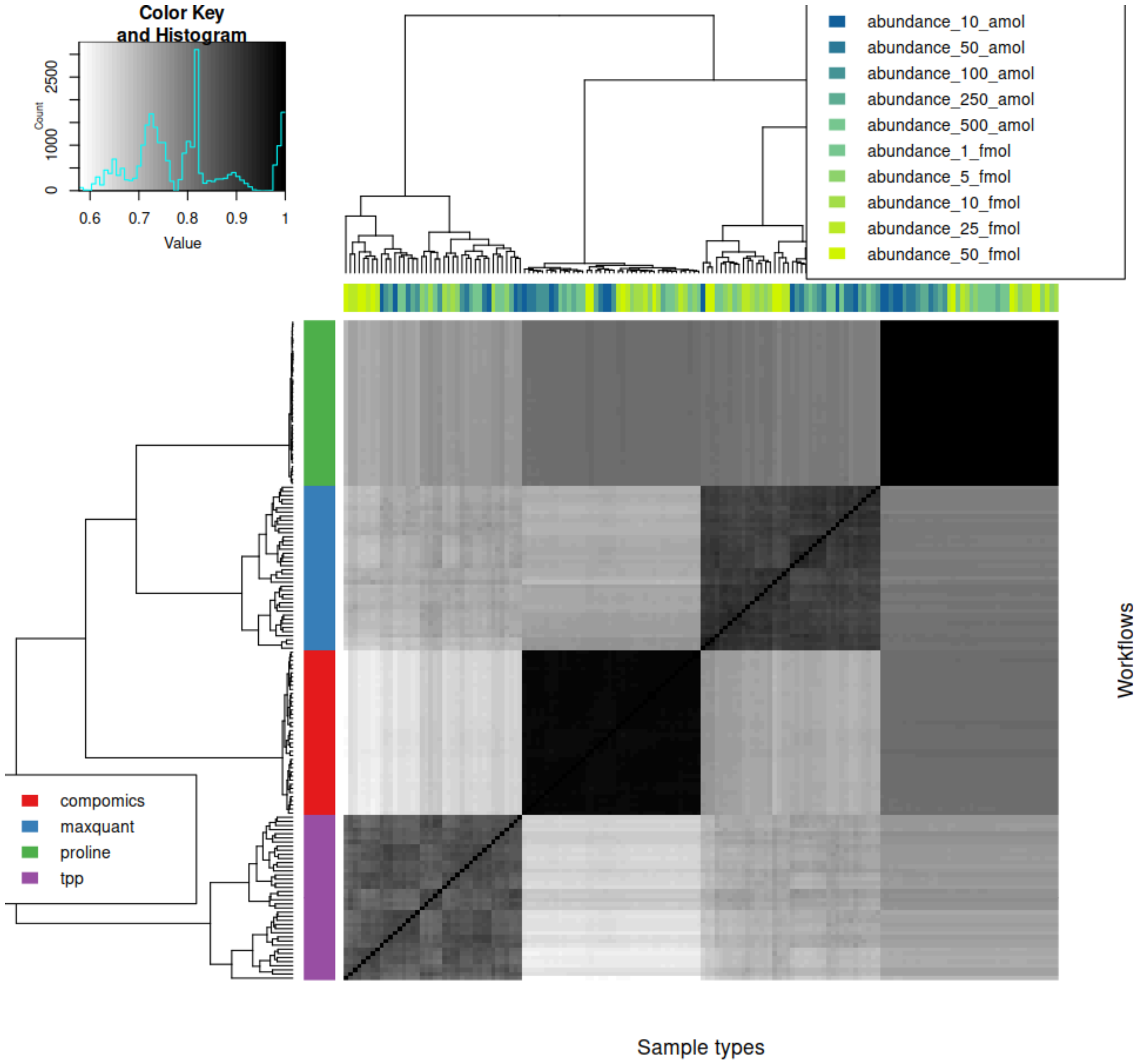


Suppl. Fig. 2: Overlap of protein identifications between all ground truth data samples analyzed by the different workflows. The similarity between identifications was calculated through Manhattan distances.

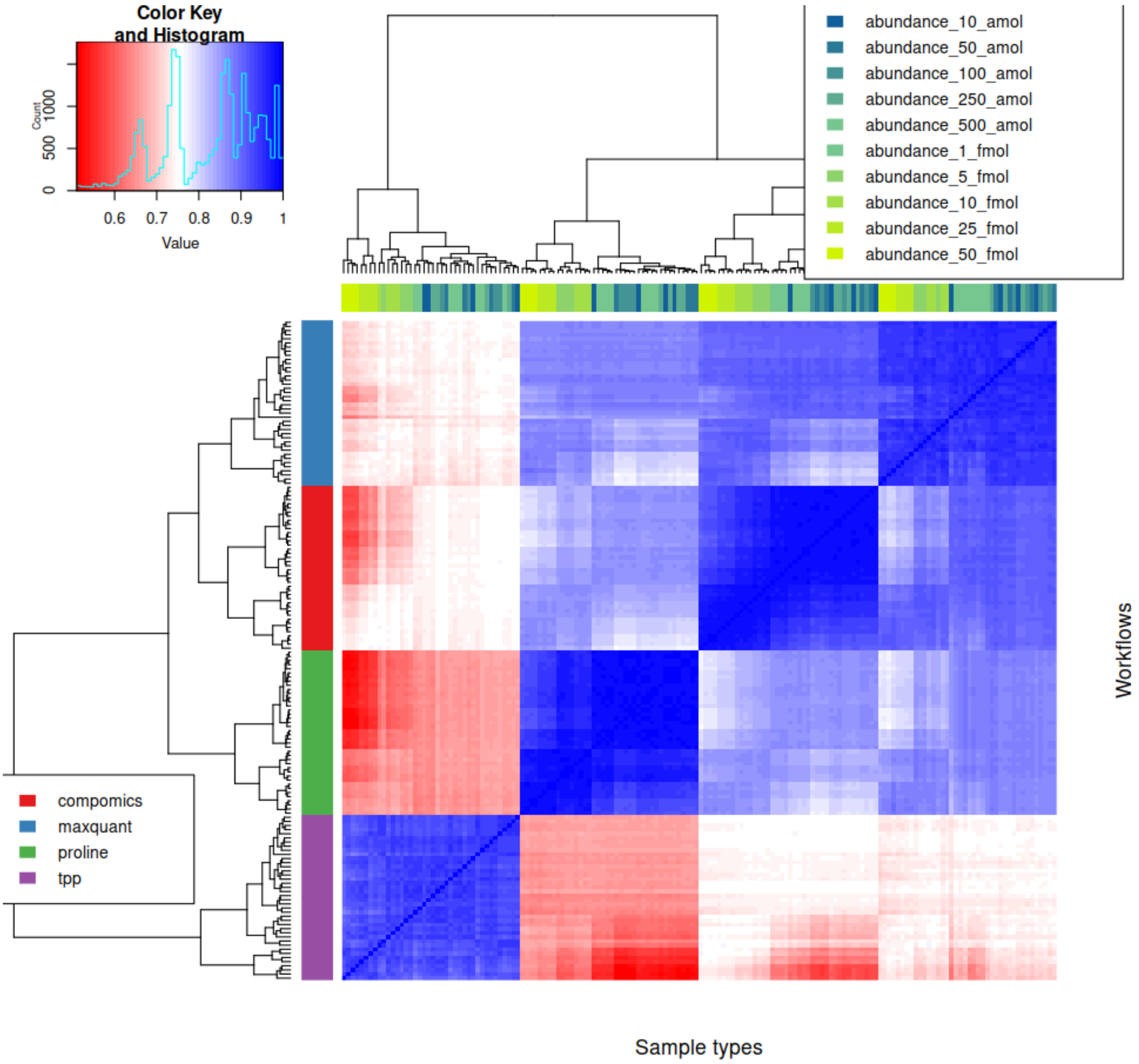


Suppl. Fig. 3: Similarity of protein quantitative profiles between different samples of the ground truth dataset (yeast spiked with UPS at different levels). We used Pearson correlation to assess the similarity. We calculated Pearson correlations to compare the similarity on a quantitative level, and clustered the correlations.

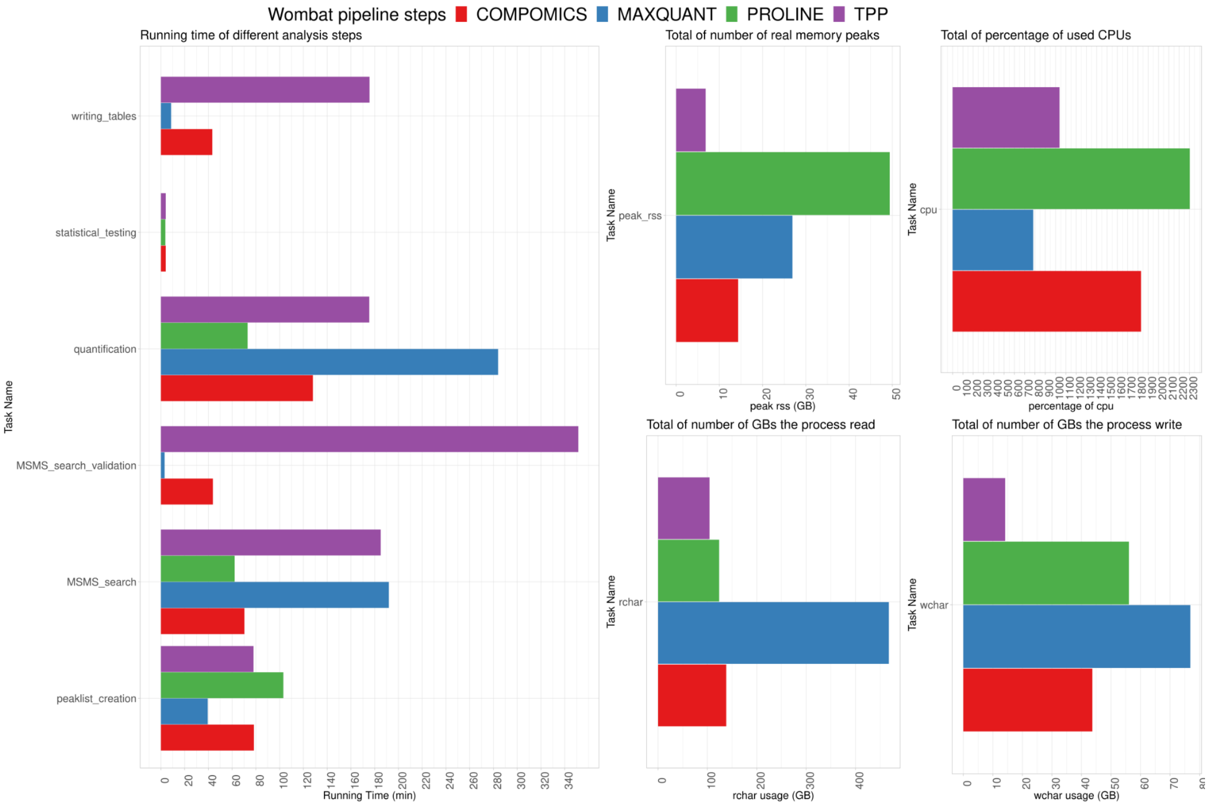


Suppl. Fig. 4: Running times for the COVID-19 dataset. rchar: amount of the data being read, wchar: amount of written data, peak_rss: peak amount of RAM, cpu: average number of used CPU threads in  percentage.

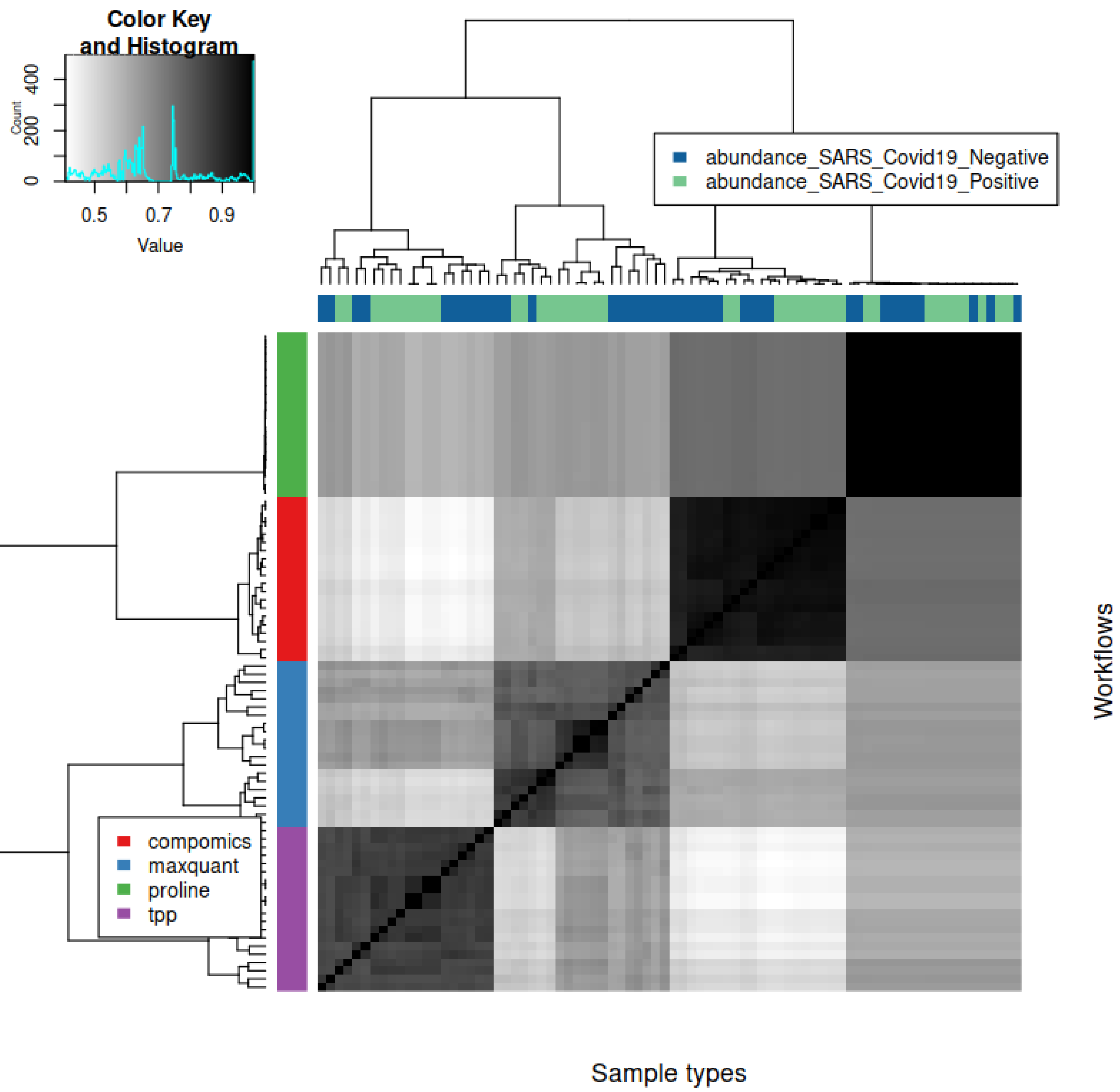


Suppl. Fig. 5: Overlap of identified proteins in the different samples and workflows for the COVID-19 dataset. We note a clear separation of the workflows. The similarity between identifications was calculated through Manhattan distances.

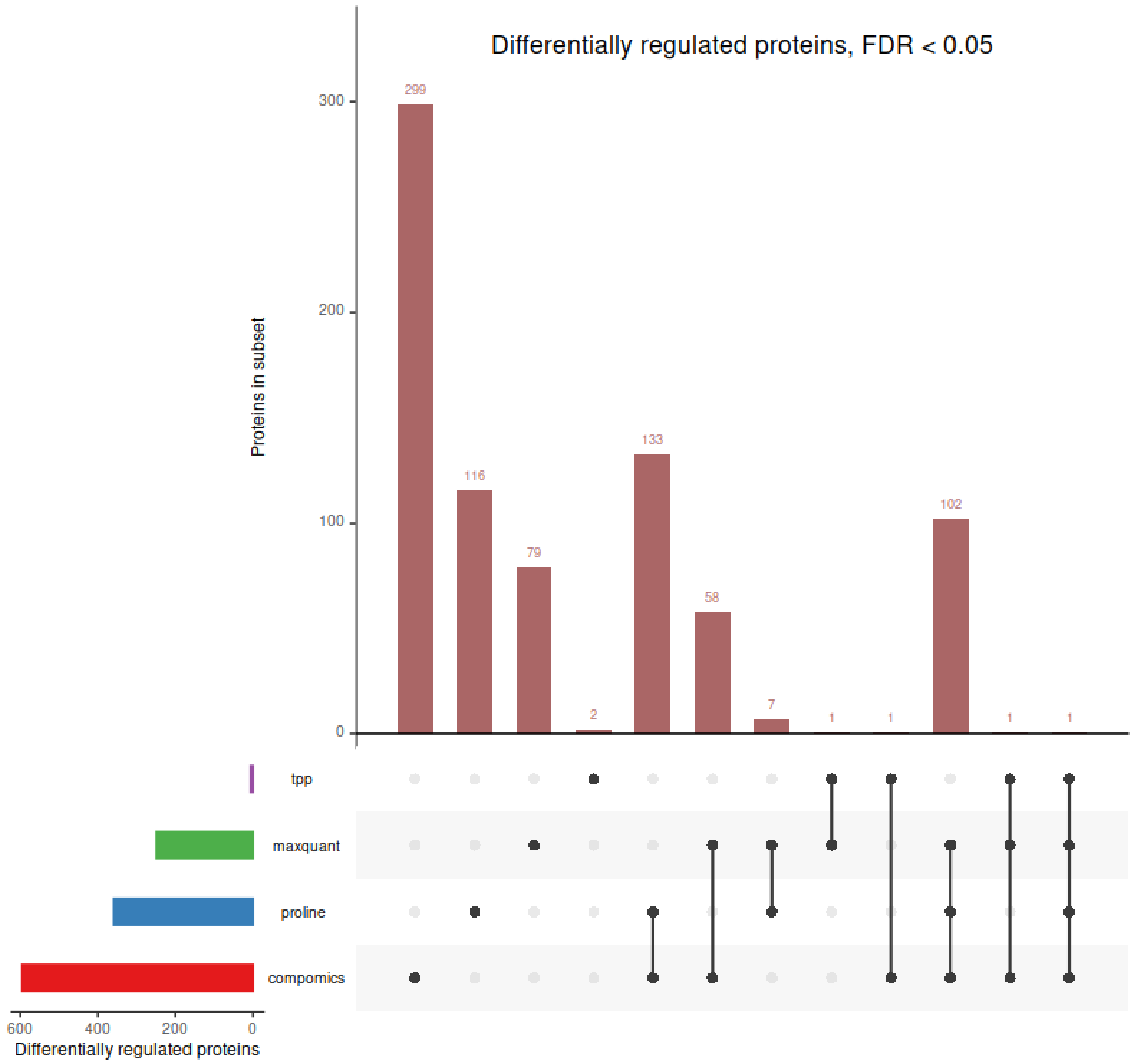


Suppl. Figure 6: Differentially regulated proteins for COVID 19 dataset for an FDR threshold of 5% showing large differences between the processing workflows.

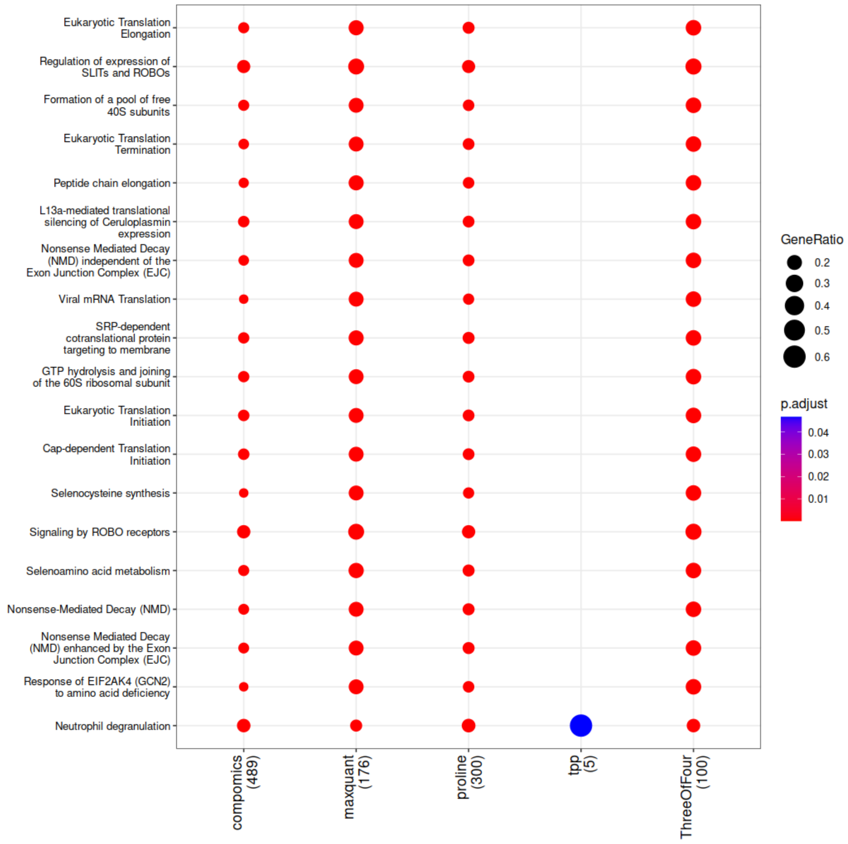


Suppl. Fig. 7: Most enriched Reactome pathways for proteins found to be differentially regulated between COVID-19 positive and negative samples (FDR for differential experssion < 0.05). Pathways are shown for the different workflows and furthermore for the proteins that were found differentially regulated in three of the four workflows. The enrichment was carried out with the clusterProfiler package.
