## Supplementary Material for "WOMBAT-P: Benchmarking Label-Free Proteomics Data Analysis Workflows"

### Wombat-P: Comparing Analysis Software in Proteomics via Heterologous Spike-in Experiments (eg UPS1)

2023-08-17

#### 1 Introduction

This document shows how **heterologous *spike-in* experiments**, like the ones using UPS1 may be analyzed in an automated manner after initial data-processing by Wombat-P.

This is script version 0.9.3.8, written by Wolfgang Raffelsberger (IGBMC, Illkirch, France)

##### 1.1 Experimental Setup For Benchmark Tests

Ground-truth experiments refer to experimental setup where the correct answer (eg which proteins do change abundance between different sets of samples) is know in advance. Heterologous *spike-in* experiments provide help for testing an comparison of identification and quantitation procedures in proteomics since the underlying ground-truth is defined by the experimental setup.

A frequent setup of spike-in experiment is based on mixing known amounts of a collection of human proteins (UPS1) in various concentrations on top of a constant level yeast total protein extract, one expects to find only the spiked human UPS1 proteins varying between samples. Sets of purified proteins have the advantage, that we know in advance which proteins *should* get detected (even if they are possibly not identified from the mass spectrometry data).

The initial data from Pride get analyzed using Wombat-P first. This workflow/script was designed specifically for experiments using heterologous *spike-in* like the ones using UPS1 (available from Sigma-Aldrich) mixed in specific ratios with yeast protein extracts as constant matrix. Anyway, the *spike-in* and matrix must always represent different protein species. If possible, using species evolutionary quite apart is preferable to avoid misclassification of peptides common to both species.

**This experiment uses UPS1 as spike-in and *Saccharomyces cerevisiae* as matrix**.

##### 1.2 Computational Setup

This workflow uses data produced by various quantification approaches integrated to Wombat-P. Then, further calculations in R make use of the additional R-packages wrProteo, wrMisc, wrGraph and RColorBrewer. All these packages are available on CRAN. Furthermore, the Bioconductor package limma will be used internally for it's moderated statistical testing.

For further details about using the R-package wrProteo and detailed examples of code, please look at the vignettes attached to this package. This workflow is derived from the UPS-1 dedicated vignette of package wrProteo.

### 2 Import Of Data Quantified In Parallel By Multiple Methods Using Wombat-P

To get started, the first step consists in importing and parsing the data. In this experiment data from **4 quantification methods** were found as previously produced by Wombat-P (see also Table 1).

During the following analysis 48 spiked UPS1 proteins are considered as 'true results' (ie 'True Positives').

Table 1: Number of proteins: by software, custom tags and species

| | Tag as spike | unknown species | Bos taurus | Homo sapiens | Mus musculus | Saccharomyces cerevisiae | Sus scrofa |
|---|---|---|---|---|---|---|---|
| compomics (CP) | 48 | 4 | 2 | 56 | 0 | 2652 | 1 |
| maxquant (MQ) | 48 | 4 | 2 | 56 | 1 | 2483 | 1 |
| proline (PL) | 48 | 4 | 2 | 58 | 3 | 2194 | 1 |
| tpp (TP) | 48 | 4 | 3 | 55 | 1 | 2499 | 0 |

#### 2.1 Normalization to *Saccharomyces cerevisiae* Matrix Proteins Only

With *spike-in* experiments it makes sense to normalize data based on the matrix proteins (here *Saccharomyces cerevisiae*), since they are experimentally kept constant. It is very beneficial to normalize only to these constant matrix-proteins since the highest concentration of spiked proteins (or peptides) may be very high and may be influencing regular global normalization approaches. Since one typically wants to measure and compare abundancess, the 'effect' of adding spiked proteins may get reduced during reguar global normalization (basic Wombat-P). By re-normalizing to the constant matrix-proteins this problem gets solved.

In consequence, at import 2 plots will be produced for each of 4 quantification methods : The first shows the distribution of protein abundance values of the initial data and the second plot the distribution after normalizing to the main species only.

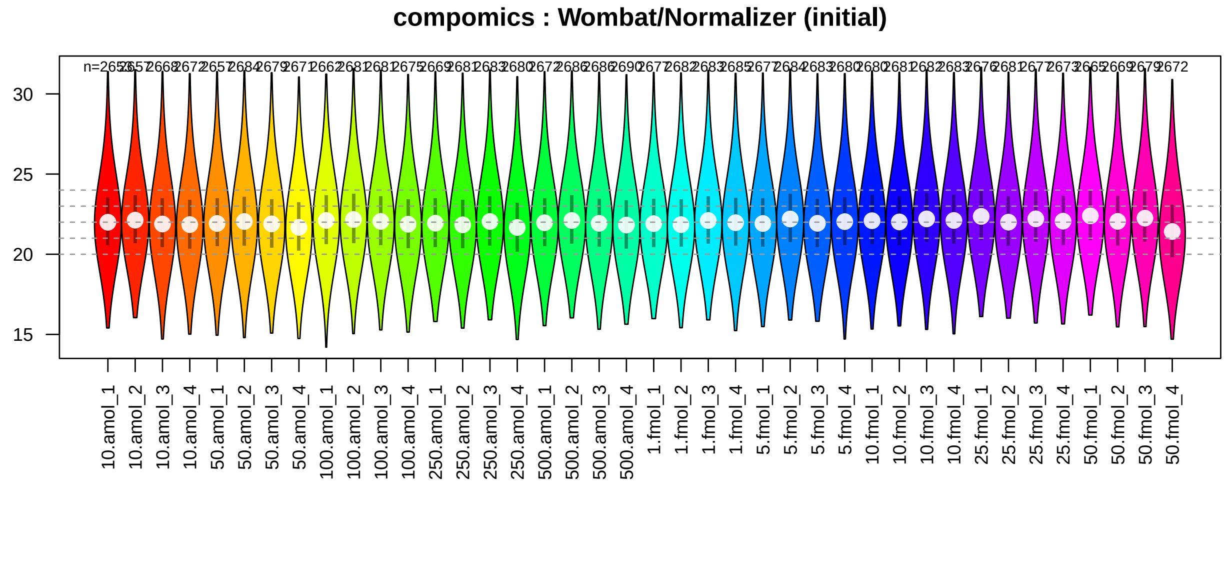

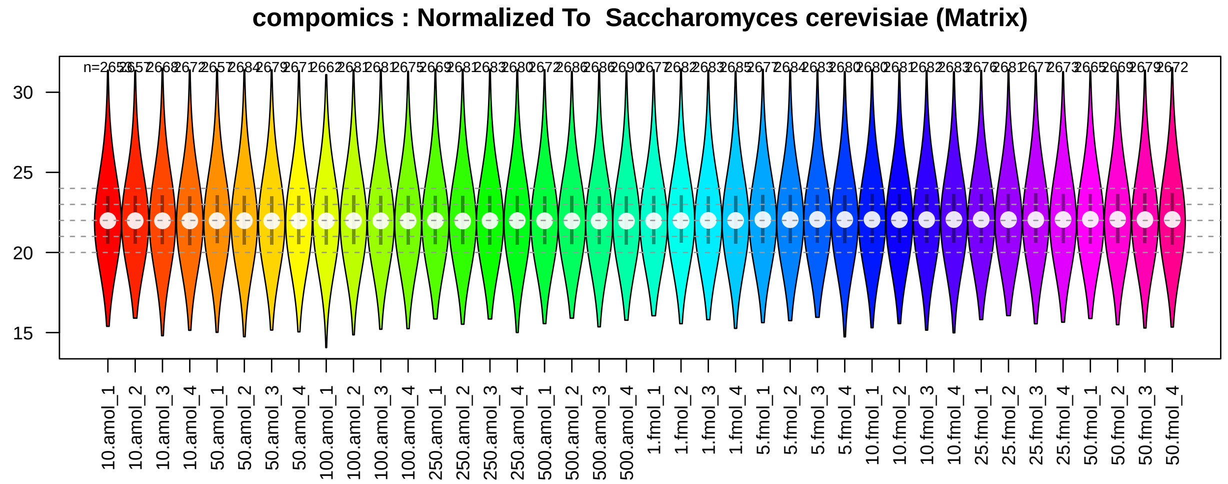
compomics : Normalized To  Saccharomyces cerevisiae (Matrix)
30
25
20
15
10.amol_1
10.amol_2
10.amol_3
10.amol_4
50.amol_1
50.amol_2
50.amol_3
50.amol_4
100.amol_1
100.amol_2
100.amol_3
100.amol_4
250.amol_1
250.amol_2
250.amol_3
250.amol_4
500.amol_1
500.amol_2
500.amol_3
500.amol_4
1.fmol_1
1.fmol_2
1.fmol_3
1.fmol_4
5.fmol_1
5.fmol_2
5.fmol_3
5.fmol_4
10.fmol_1
10.fmol_2
10.fmol_3
10.fmol_4
25.fmol_1
25.fmol_2
25.fmol_3
25.fmol_4
50.fmol_1
50.fmol_2
50.fmol_3
50.fmol_4

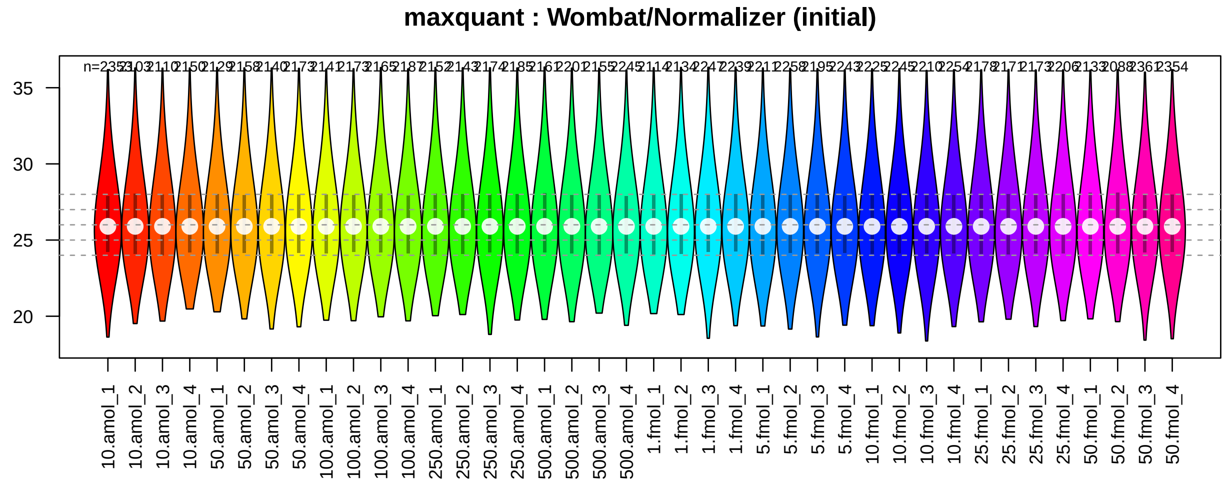
maxquant : Wombat/Normalizer (initial)
35
30
25
20
10.amol_1
10.amol_2
10.amol_3
10.amol_4
50.amol_1
50.amol_2
50.amol_3
50.amol_4
100.amol_1
100.amol_2
100.amol_3
100.amol_4
250.amol_1
250.amol_2
250.amol_3
250.amol_4
500.amol_1
500.amol_2
500.amol_3
500.amol_4
1.fmol_1
1.fmol_2
1.fmol_3
1.fmol_4
5.fmol_1
5.fmol_2
5.fmol_3
5.fmol_4
10.fmol_1
10.fmol_2
10.fmol_3
10.fmol_4
25.fmol_1
25.fmol_2
25.fmol_3
25.fmol_4
50.fmol_1
50.fmol_2
50.fmol_3
50.fmol_4

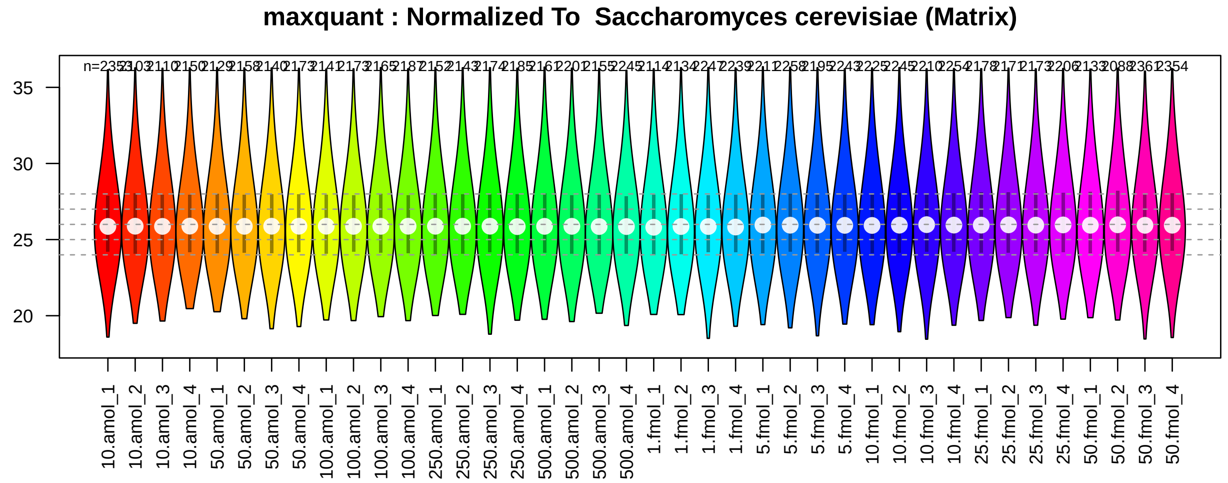
maxquant : Normalized To  Saccharomyces cerevisiae (Matrix)
35
30
25
20
10.amol_1
10.amol_2
10.amol_3
10.amol_4
50.amol_1
50.amol_2
50.amol_3
50.amol_4
100.amol_1
100.amol_2
100.amol_3
100.amol_4
250.amol_1
250.amol_2
250.amol_3
250.amol_4
500.amol_1
500.amol_2
500.amol_3
500.amol_4
1.fmol_1
1.fmol_2
1.fmol_3
1.fmol_4
5.fmol_1
5.fmol_2
5.fmol_3
5.fmol_4
10.fmol_1
10.fmol_2
10.fmol_3
10.fmol_4
25.fmol_1
25.fmol_2
25.fmol_3
25.fmol_4
50.fmol_1
50.fmol_2
50.fmol_3
50.fmol_4

**proline : Wombat/Normalizer (initial)**

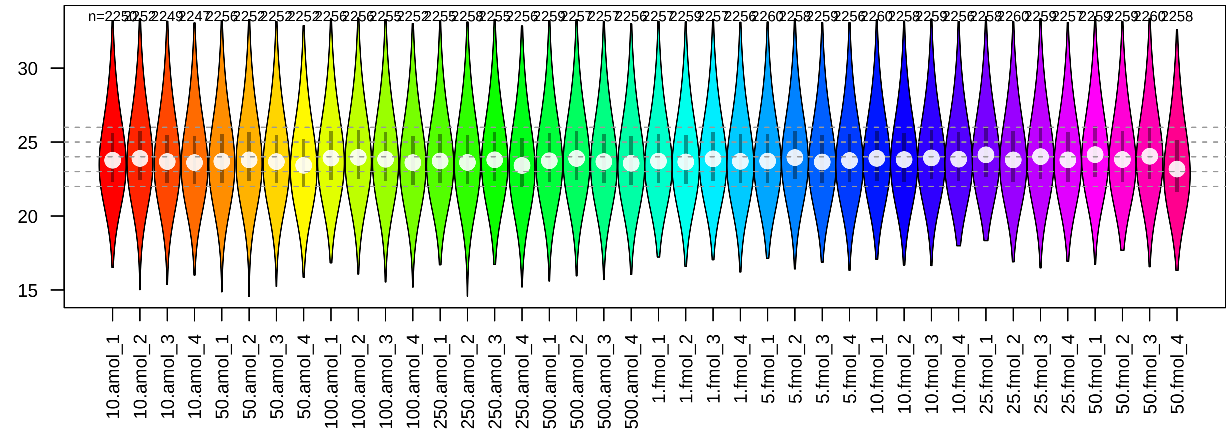


**proline : Normalized To Saccharomyces cerevisiae (Matrix)**

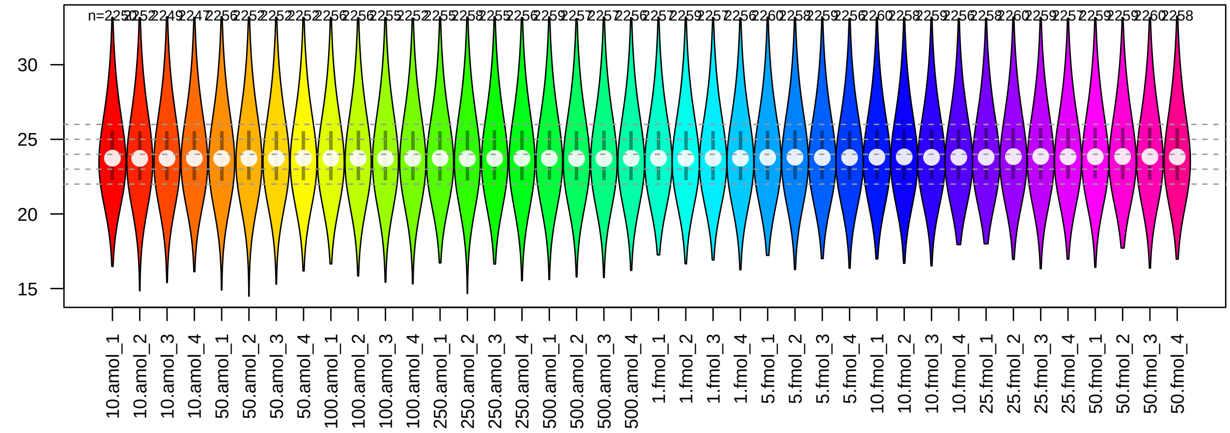


**tpp : Wombat/Normalizer (initial)**

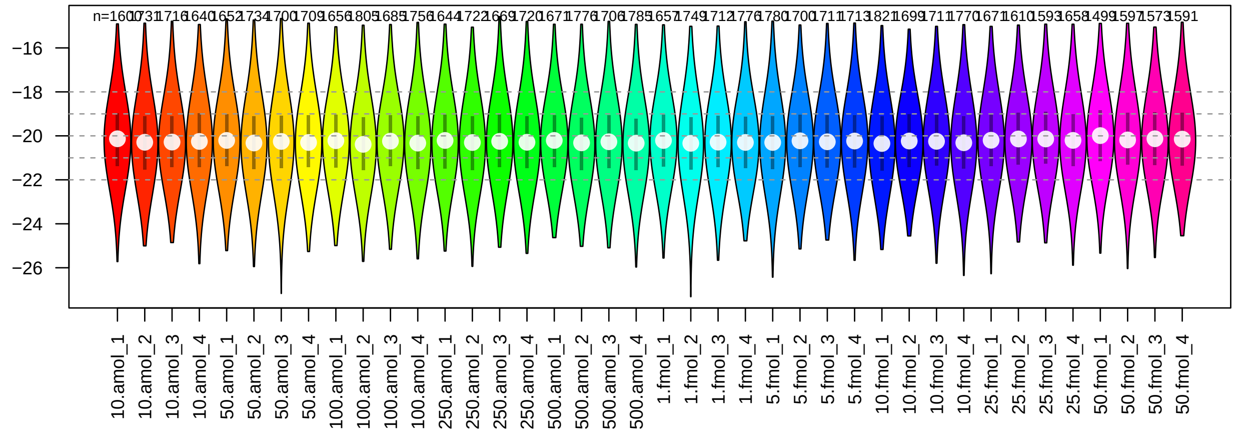

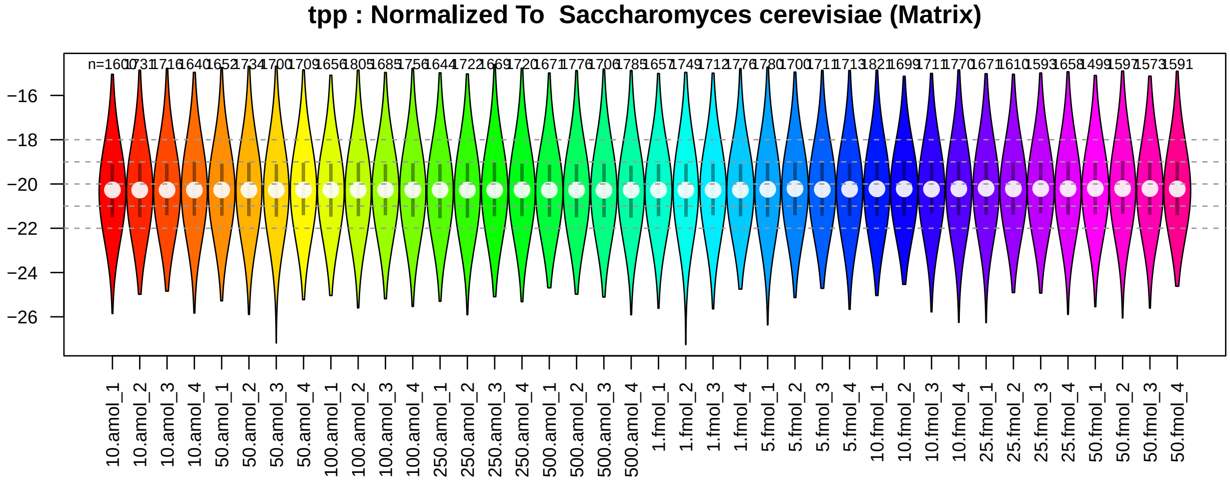


#### 2.2 Inspecting and Addressing NA-Values

It is important to investigate the nature of NA-values, as explained in the general 'wrProteoVignette1' of the package wrProteo. In particular, checking the hypothesis that NA-values originate from very low abundance instances is very important for deciding how to treat NA-values furtheron. The figure below shows the distribution of all data (grey) and NA-Neighbours (green and blue).

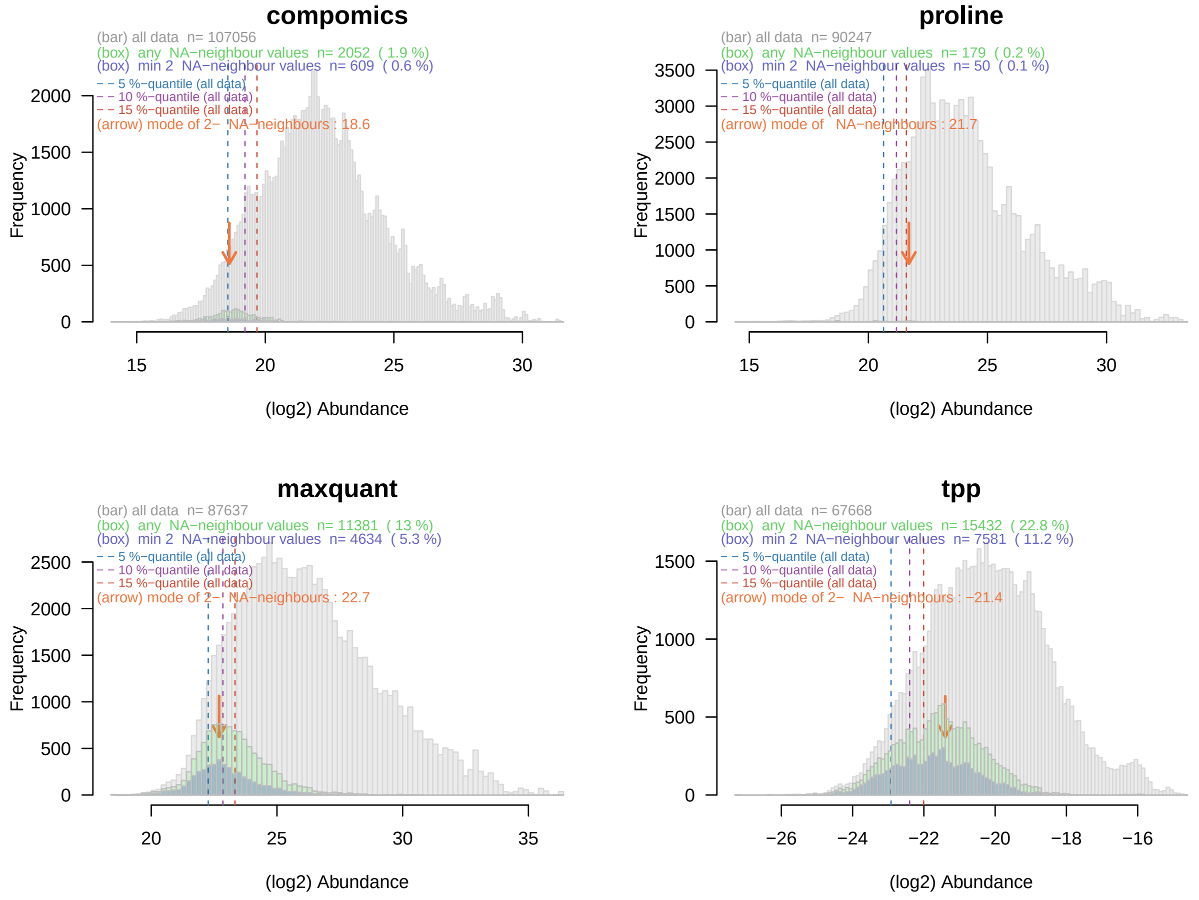

##### 2.2.1 NA-Imputation and Statistical Testing for Changes in Abundance

NA-values represent a challange for statistical testing. In addition, techniques like PCA don't allow NAs, neither.

The number of NAs varies between quantitation methods and between samples : Indeed, very low concentrations of spike-in proteins (UPS1) are difficult to get detected and contribute to the NAs (as we will see later in more detail). Since the amout of main matrix proteins (*Saccharomyces cerevisiae*) stays constant across all samples, matrix proteins should always get detected the same way. NA-imputation and testing for quantification data is performed as described in wrProteo.

Table 2: Number of NAs per group of samples

| | 10amol | 50amol | 100amol | 250amol | 500amol | 1000amol | 5000amol | 10000amol | 25000amol | 50000amol |
|---|---|---|---|---|---|---|---|---|---|---|
| compomics | 210 | 169 | 161 | 147 | 126 | 133 | 136 | 134 | 153 | 175 |
| maxquant | 1484 | 1600 | 1534 | 1546 | 1438 | 1466 | 1293 | 1266 | 1472 | 1264 |
| proline | 50 | 36 | 29 | 24 | 19 | 19 | 15 | 15 | 14 | 12 |
| tpp | 3561 | 3453 | 3346 | 3493 | 3310 | 3354 | 3344 | 3247 | 3716 | 3988 |

##### 2.2.2 CV of Replicates

As general indicator for data quality and usability let's look at intra-replicate variability. First we calculate for each protein and set of replicate-samples the corresponding CV. Then, we can plot the distribution of all CV values as violin plots which show a kernel-estimate for the distribution. In addition, a box-plot is also integrated to each distribution (see also vignette to package wrGraph for details).

In the figure below the distributions oriented to left side on figure corresponds to the CV values from all proteins, and the ones oriented to right side only from the spiked proteins. You can see that low and high amounts of spiked proteins have quite different CVs.

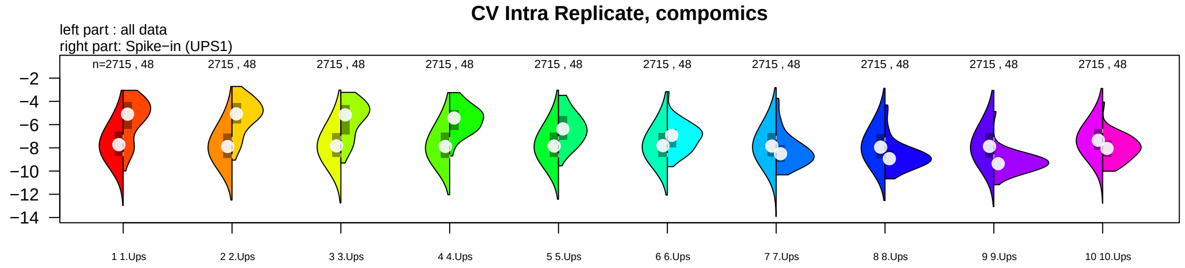


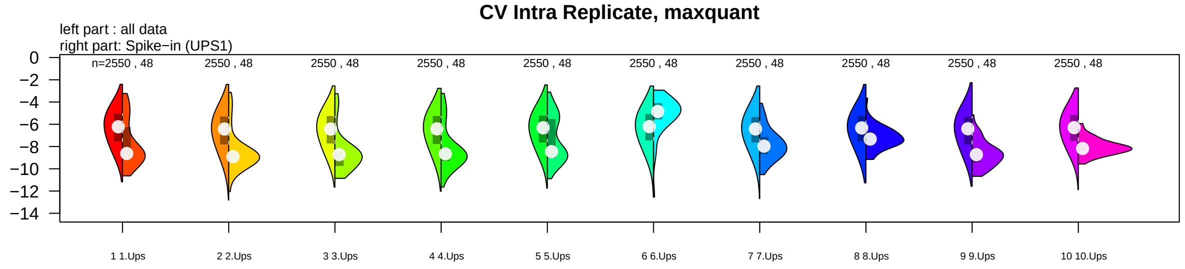

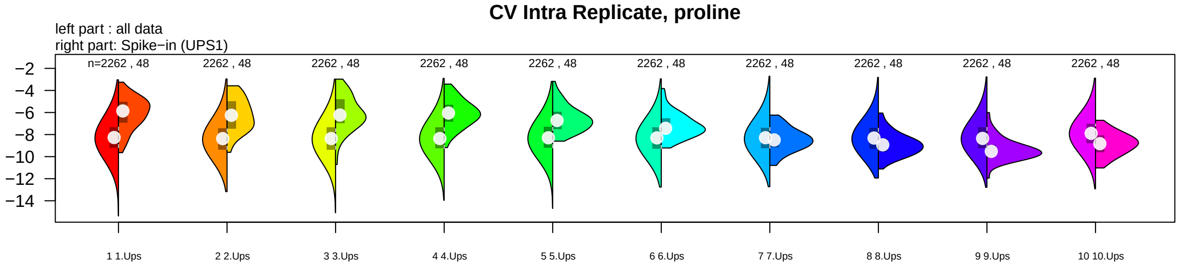


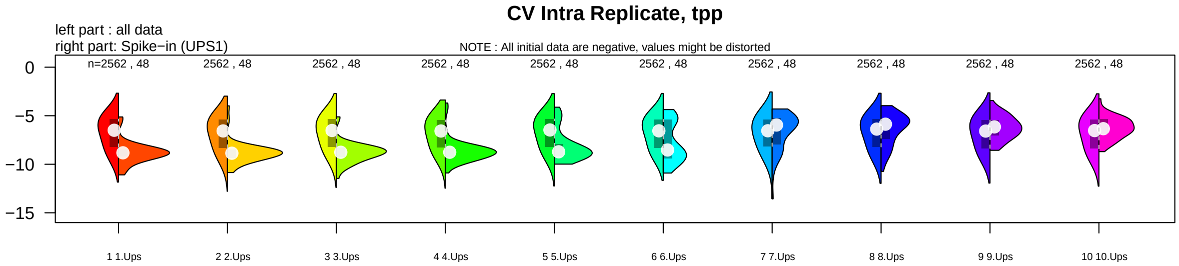


### 3 Analysis Based On Pair-Wise Testing Of All Proteins (Matrix And Spike-In)

In this section we consider all proteins identified and quantified in a pair-wise fashion using t-tests. As mentioned, the experimental setup in *spike-in* experiments is very special, since all proteins that are truly changing abundance are known in advance while matrix proteins should remain constant.

For each pair-wise comparison all proteins observed as changing abundance above a traditional 5 percent Benjamini-Hochberg FDR cut-off are retained as 'differentially abundant'. Then, tables giving the number of True Positives (TP, ie the number of spiked proteins expected to change and indeed found differential in the experimental data) can be constructed. Single pair-wise comparisons can be shown as Volcano-plots (using a 5 percent FDR cut-off), while ROC-curves allow inspecting the entire range of potential cut-off values.

#### 3.1 Pairwise Testing Summary

Now we inspect Table 3 below, showing all possible pairwise-comparisons. It displays the (log-)ratio of the pairwise comparisons (column 'log2rat'), the concentrations of spiked proteins (columns 'conc1' and 'conc2', brought to a common unit-denominator) and of course, the number of differentially abundant proteins passing 5% Benjamini-Hochberg FDR.

Table 3: All pairwise comparisons and number of significant proteins

| | index | log2rat | conc1 | conc2 | nCP.BH | nMQ.BH | nPL.BH | nTP.BH |
|---|---|---|---|---|---|---|---|---|
| 10amol-50000amol | 19 | 12.290 | 10 | 50000 | 233 | 26 | 259 | 14 |
| 10amol-25000amol | 10 | 11.290 | 10 | 25000 | 71 | 22 | 110 | 4 |
| 10000amol-10amol | 4 | 9.966 | 10 | 10000 | 52 | 3 | 75 | 0 |
| 50000amol-50amol | 43 | 9.966 | 50 | 50000 | 214 | 66 | 221 | 62 |
| 100amol-50000amol | 18 | 8.966 | 100 | 50000 | 212 | 27 | 249 | 42 |
| 10amol-5000amol | 25 | 8.966 | 10 | 5000 | 46 | 4 | 51 | 1 |

| | index | log2rat | conc1 | conc2 | nCP.BH | nMQ.BH | nPL.BH | nTP.BH |
|---|---|---|---|---|---|---|---|---|
| 25000amol-50amol | 41 | 8.966 | 50 | 25000 | 76 | 37 | 115 | 9 |
| 100amol-25000amol | 9 | 7.966 | 100 | 25000 | 64 | 26 | 94 | 3 |
| 250amol-50000amol | 21 | 7.644 | 250 | 50000 | 228 | 37 | 245 | 22 |
| 10000amol-50amol | 37 | 7.644 | 50 | 10000 | 49 | 4 | 55 | 1 |
| 10000amol-100amol | 2 | 6.644 | 100 | 10000 | 46 | 4 | 52 | 0 |
| 1000amol-10amol | 5 | 6.644 | 10 | 1000 | 31 | 3 | 26 | 0 |
| 25000amol-250amol | 15 | 6.644 | 250 | 25000 | 74 | 38 | 105 | 13 |
| 50000amol-500amol | 35 | 6.644 | 500 | 50000 | 192 | 84 | 230 | 89 |
| 5000amol-50amol | 44 | 6.644 | 50 | 5000 | 51 | 55 | 54 | 30 |
| 1000amol-50000amol | 17 | 5.644 | 1000 | 50000 | 192 | 25 | 240 | 29 |
| 100amol-5000amol | 24 | 5.644 | 100 | 5000 | 40 | 3 | 49 | 0 |
| 10amol-500amol | 32 | 5.644 | 10 | 500 | 23 | 0 | 27 | 0 |
| 25000amol-500amol | 33 | 5.644 | 500 | 25000 | 62 | 27 | 107 | 11 |
| 10000amol-250amol | 11 | 5.322 | 250 | 10000 | 49 | 5 | 59 | 2 |
| 1000amol-25000amol | 8 | 4.644 | 1000 | 25000 | 59 | 14 | 103 | 6 |
| 10amol-250amol | 14 | 4.644 | 10 | 250 | 14 | 0 | 8 | 0 |
| 250amol-5000amol | 27 | 4.322 | 250 | 5000 | 47 | 46 | 50 | 16 |
| 10000amol-500amol | 29 | 4.322 | 500 | 10000 | 46 | 5 | 55 | 1 |
| 1000amol-50amol | 38 | 4.322 | 50 | 1000 | 36 | 3 | 34 | 0 |
| 10000amol-1000amol | 1 | 3.322 | 1000 | 10000 | 44 | 4 | 57 | 1 |
| 1000amol-100amol | 3 | 3.322 | 100 | 1000 | 30 | 3 | 28 | 0 |
| 100amol-10amol | 6 | 3.322 | 10 | 100 | 5 | 0 | 0 | 0 |
| 50000amol-5000amol | 28 | 3.322 | 5000 | 50000 | 166 | 75 | 197 | 76 |
| 5000amol-500amol | 36 | 3.322 | 500 | 5000 | 46 | 50 | 52 | 28 |
| 500amol-50amol | 45 | 3.322 | 50 | 500 | 22 | 3 | 28 | 0 |
| 10000amol-50000amol | 16 | 2.322 | 10000 | 50000 | 104 | 14 | 139 | 17 |
| 1000amol-5000amol | 23 | 2.322 | 1000 | 5000 | 37 | 2 | 43 | 1 |
| 25000amol-5000amol | 26 | 2.322 | 5000 | 25000 | 55 | 23 | 94 | 7 |
| 100amol-500amol | 31 | 2.322 | 100 | 500 | 14 | 2 | 22 | 0 |
| 10amol-50amol | 40 | 2.322 | 10 | 50 | 5 | 0 | 10 | 0 |
| 250amol-50amol | 42 | 2.322 | 50 | 250 | 12 | 0 | 9 | 0 |
| 1000amol-250amol | 12 | 2.000 | 250 | 1000 | 29 | 2 | 25 | 0 |
| 10000amol-25000amol | 7 | 1.322 | 10000 | 25000 | 27 | 0 | 56 | 3 |
| 100amol-250amol | 13 | 1.322 | 100 | 250 | 6 | 0 | 5 | 0 |
| 25000amol-50000amol | 20 | 1.000 | 25000 | 50000 | 52 | 1 | 89 | 0 |
| 10000amol-5000amol | 22 | 1.000 | 5000 | 10000 | 18 | 0 | 39 | 0 |
| 1000amol-500amol | 30 | 1.000 | 500 | 1000 | 9 | 2 | 11 | 0 |
| 250amol-500amol | 34 | 1.000 | 250 | 500 | 3 | 2 | 7 | 0 |
| 100amol-50amol | 39 | 1.000 | 50 | 100 | 3 | 0 | 1 | 0 |

You can see that in numerous cases many more than the 48 spike-in proteins (UPS1 proteins) showed up significantly, ie *Saccharomyces cerevisiae* proteins supposed to remain constant also showed up in part as 'sigificantly changing'. However, some proteins with enthousiastic FDR values have very low log-FC amplitude and will be removed by filtering in the following steps. The figure below uses Benjmini-Hochberg (BH) FDR values.

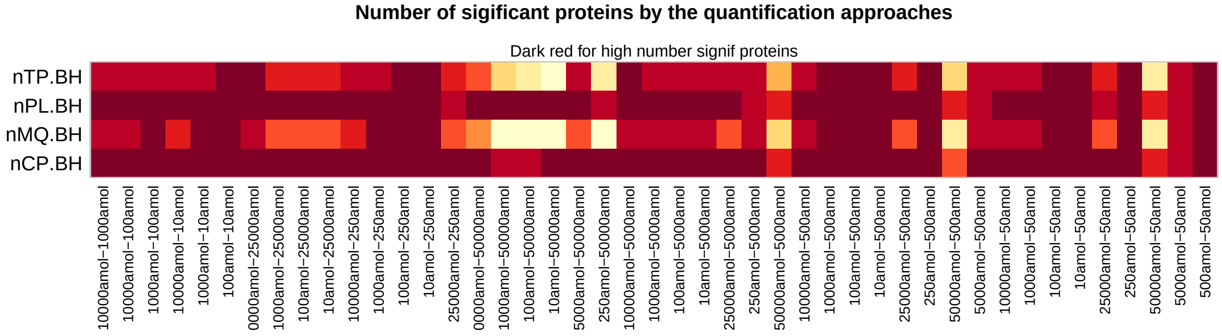

##### 3.1.1 Volcano Plots

Volcano-plots offer additional insight in how statistical test results relate to log-fold-change of pair-wise comparisons, see also vignettes to the R-packages wrProteo and wrGraph. Since the number of possible pair-wise combinations is quite elevated, only some *representative* Volcano-plots will be shown in the later sections.

##### 3.1.2 ROC for Multiple Pairs

*Receiver Operator Curves* (ROC) curves display *sensitivity* (True Positive Rate) versus *1-Specificity* (False Positive Rate), see also also the original publication Hand and Till 2001 or the vignettes to the R-package wrProteo. A good performance indicator is then the area below each of these curves (AUC).

Please note, that if some software identifies too few spike-in proteins (UPS1) for calculating sensitivity and accuracy, this may/will pose limitations later on.

#### 3.2 Gruping Of Pair-Wise Testing Results

As mentioned, there are too many pair-wise combinations available for plotting and inspecting all ROC-curves. For this reason the data will be grouped automatically in 5 sets (similar to restaurant ratings) : *best / good / average / weak / weakest* . Details of this procedure are also described in the UPS-1 dedicated vignette. Then, a single representative pair-wise comparison will be displayed for the groups '*best*', '*average*' and '*weakest*'.

Of course each of the 4 quantification methods gives different results and in consequence different AUC values (summarizing each ROC curve).

To graphically summarize the AUC values, the grouped AUC values are plotted accompanied by the geometric mean summarizing across the 4 quantification methods. In the figure below the first group from the left (ie '*best*', labelled as 'clu 5') contains the pair-wise comparisons with the best results. This means, that in this group *spike-in* proteins were frequently found as differential while matrix proteins were mainly found as not differential.

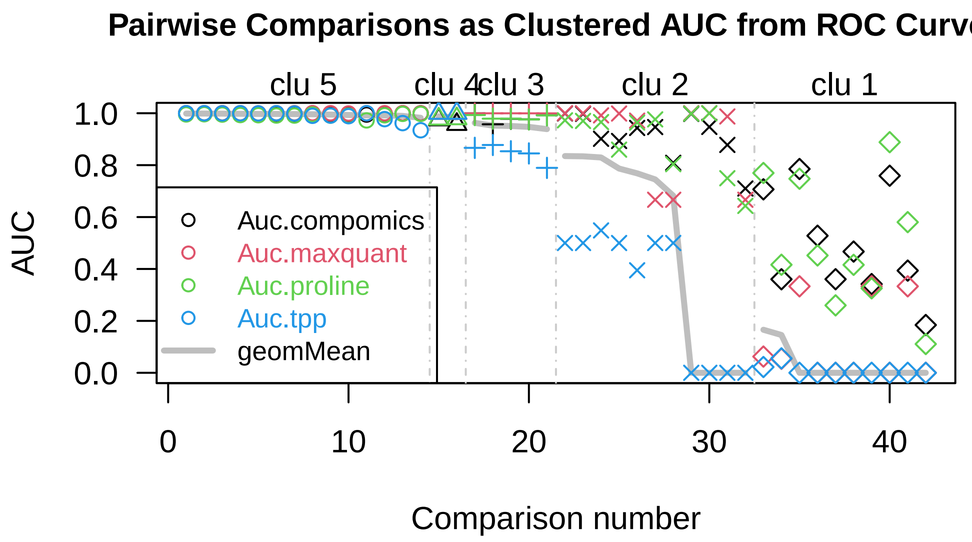


From this figure we can see clearly that there are some pairwise comparisons where all analysis-software results yield high AUC values, while other pairwise comparisons have less discriminative power between spiked and matrix-proteins.

##### 3.2.1 Plotting ROC Curves for the Best Group (the '+++++')

The best group contains 14 pair-wise comparisons shown in the table underneith.

Table 4: AUC details for the group of best pairwise-comparisons

| | Auc.CP | Auc.MQ | Auc.PL | Auc.TP | nCP.BH | nMQ.BH | nPL.BH | nTP.BH |
|---|---|---|---|---|---|---|---|---|
| 10amol-25000amol | 0.999 | 1.000 | 0.995 | 1.000 | 71 | 22 | 110 | 4 |
| 10000amol-50amol | 0.999 | 1.000 | 0.996 | 1.000 | 49 | 4 | 55 | 1 |
| 25000amol-250amol | 0.999 | 1.000 | 0.995 | 1.000 | 74 | 38 | 105 | 13 |
| 50000amol-50amol | 0.998 | 1.000 | 0.993 | 1.000 | 214 | 66 | 221 | 62 |
| 50000amol-500amol | 0.998 | 0.999 | 0.992 | 0.999 | 192 | 84 | 230 | 89 |
| 10amol-50000amol | 0.998 | 1.000 | 0.991 | 1.000 | 233 | 26 | 259 | 14 |
| 250amol-50000amol | 0.997 | 1.000 | 0.991 | 0.999 | 228 | 37 | 245 | 22 |
| 1000amol-25000amol | 0.999 | 1.000 | 0.995 | 0.990 | 59 | 14 | 103 | 6 |
| 1000amol-50000amol | 0.997 | 1.000 | 0.991 | 0.991 | 192 | 25 | 240 | 29 |
| 50000amol-5000amol | 0.995 | 0.999 | 0.990 | 0.989 | 166 | 75 | 197 | 76 |
| 10000amol-250amol | 0.995 | 1.000 | 0.972 | 1.000 | 49 | 5 | 59 | 2 |
| 100amol-50000amol | 0.998 | 1.000 | 0.992 | 0.977 | 212 | 27 | 249 | 42 |
| 100amol-25000amol | 1.000 | 1.000 | 0.998 | 0.962 | 64 | 26 | 94 | 3 |
| 25000amol-50amol | 1.000 | 0.999 | 0.998 | 0.934 | 76 | 37 | 115 | 9 |

Then we can see which spike-in amounts/concentrations were found in these pair-wise comparisons (histogram below), display the ROC-curves for each of the 4 analysis-software and finally the corresponding Volcano-plots.

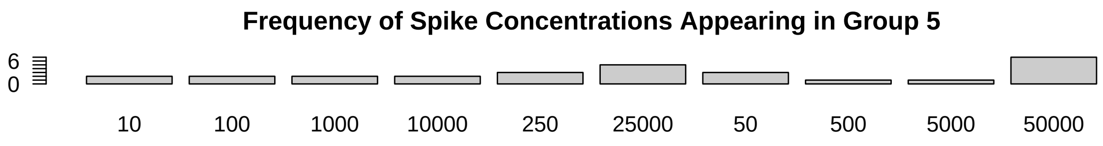


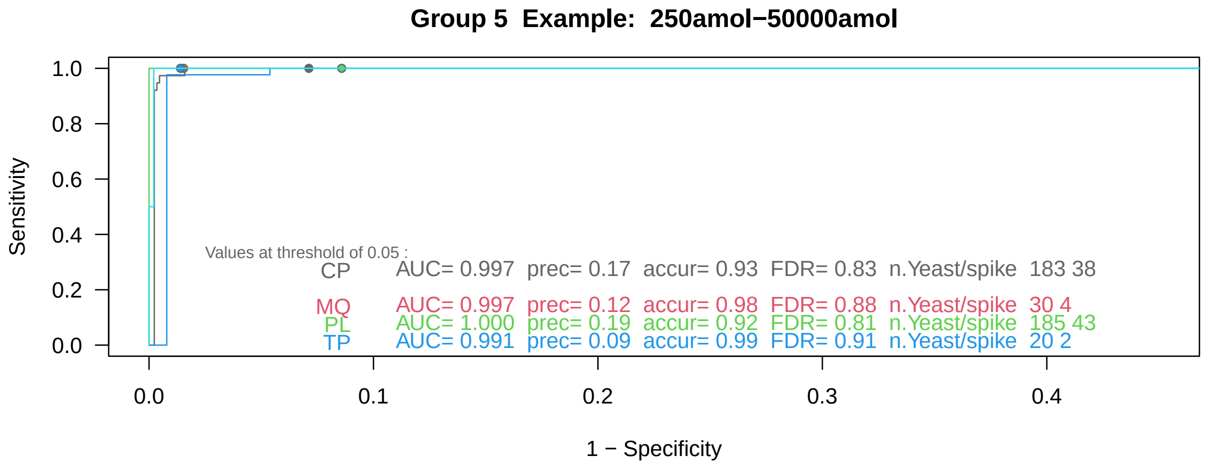

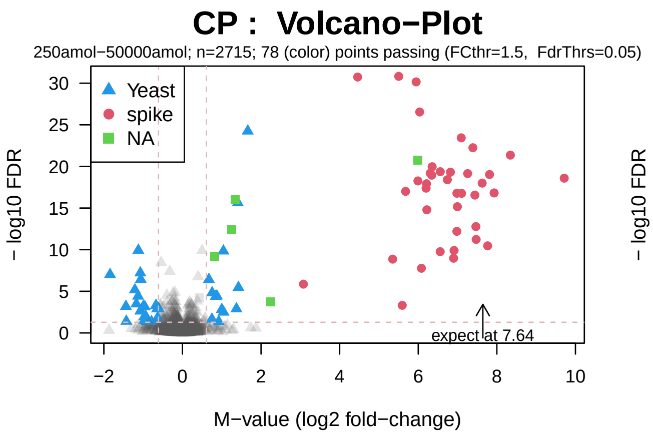


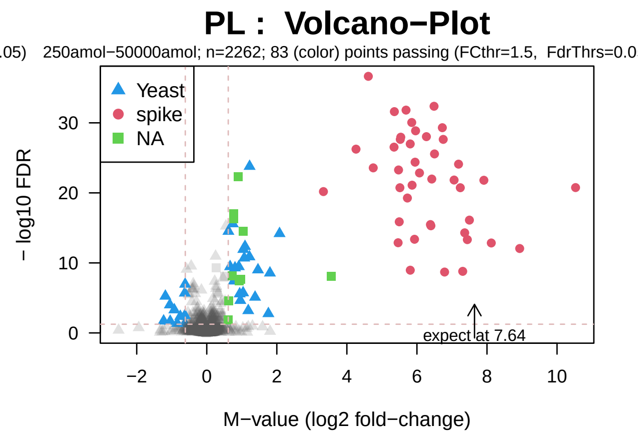


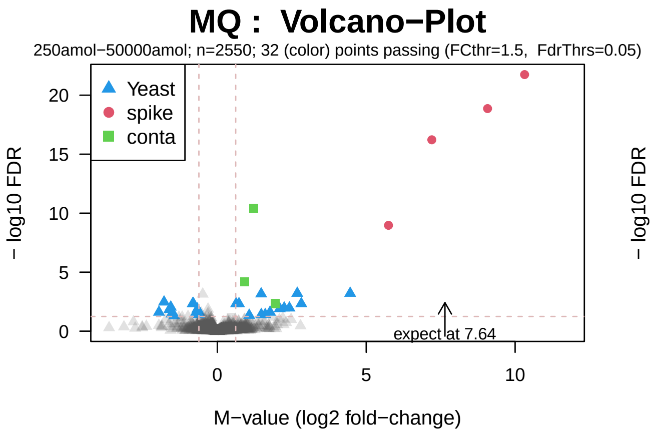


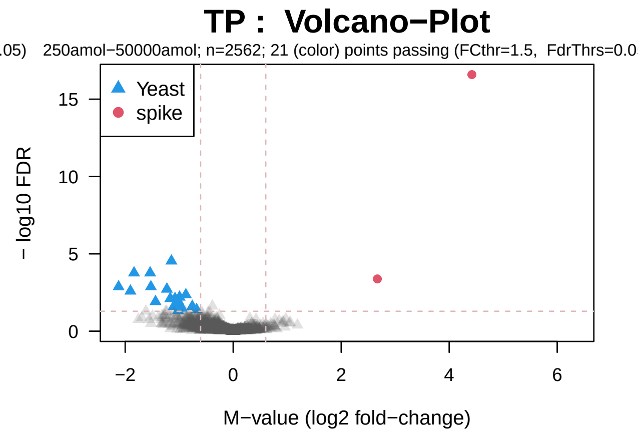


##### 3.2.2 ROC Curves for the 'Average' Group (the '+++')

The 'average' group contains 5 pair-wise comparisons shown in the table underneith.

Table 5: AUC details for best pairwise-comparisons

| | Auc.CP | Auc.MQ | Auc.PL | Auc.TP | nCP.BH | nMQ.BH | nPL.BH | nTP.BH |
|---|---|---|---|---|---|---|---|---|
| 25000amol-500amol | 1.000 | 0.999 | 0.993 | 0.867 | 62 | 27 | 107 | 11 |
| 5000amol-50amol | 0.957 | 1.000 | 0.979 | 0.878 | 51 | 55 | 54 | 30 |
| 5000amol-500amol | 0.978 | 1.000 | 0.979 | 0.853 | 46 | 50 | 52 | 28 |
| 25000amol-5000amol | 0.977 | 1.000 | 0.976 | 0.845 | 55 | 23 | 94 | 7 |
| 10000amol-50000amol | 0.992 | 0.999 | 0.992 | 0.789 | 104 | 14 | 139 | 17 |

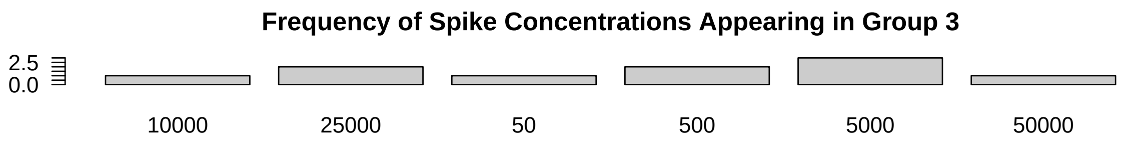
Frequency of Spike Concentrations Appearing in Group 3
2.5
0.0
10000
25000
50
500
5000
50000

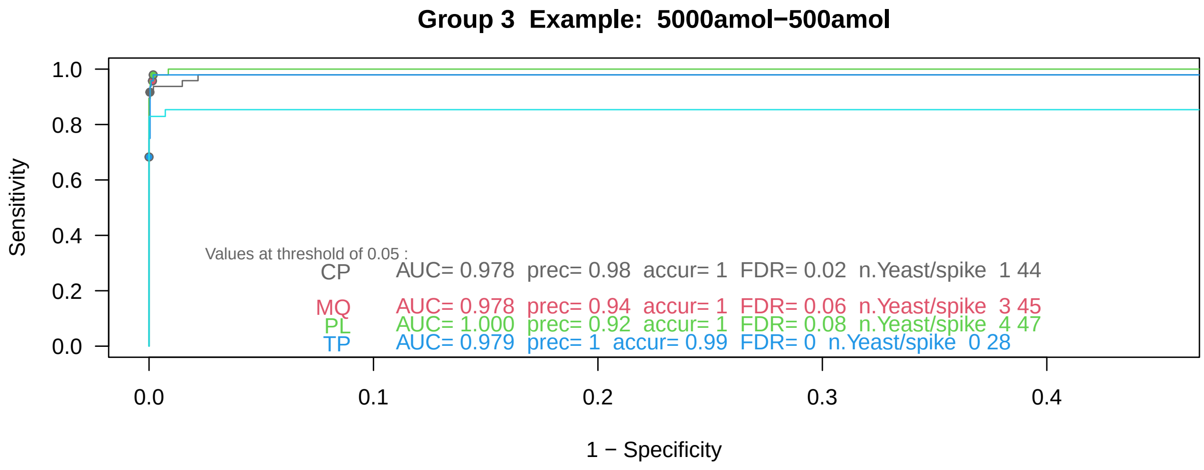
Group 3 Example: 5000amol−500amol
Sensitivity
1 − Specificity
Values at threshold of 0.05 :
CP AUC= 0.978 prec= 0.98 accur= 1 FDR= 0.02 n.Yeast/spike 1 44
MQ AUC= 0.978 prec= 0.94 accur= 1 FDR= 0.06 n.Yeast/spike 3 45
PL AUC= 1.000 prec= 0.92 accur= 1 FDR= 0.08 n.Yeast/spike 4 47
TP AUC= 0.979 prec= 1 accur= 0.99 FDR= 0 n.Yeast/spike 0 28

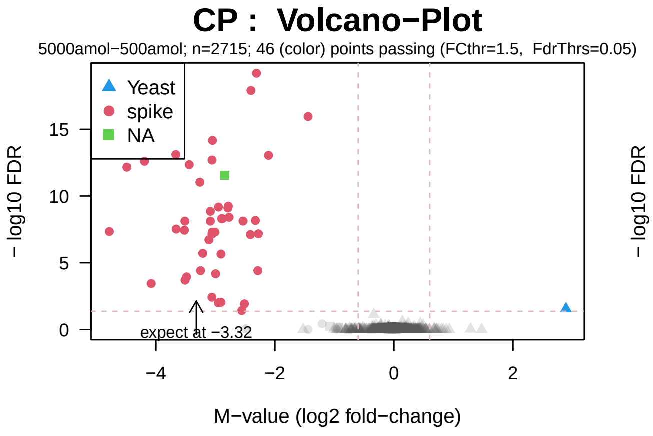
CP : Volcano−Plot
5000amol−500amol; n=2715; 46 (color) points passing (FCthr=1.5, FdrThrs=0.05)
Yeast
spike
NA
expect at −3.32
− log10 FDR
M−value (log2 fold−change)

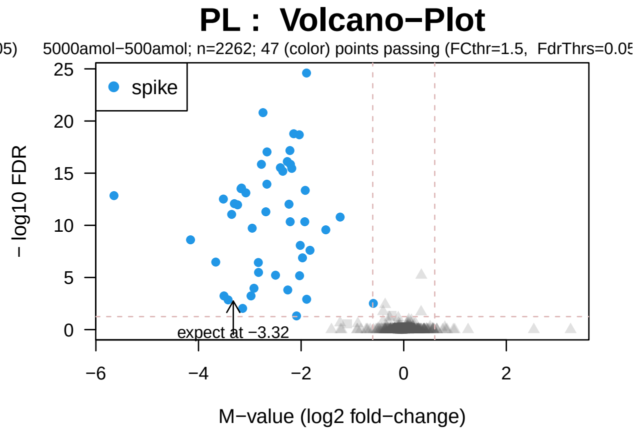
PL : Volcano−Plot
5000amol−500amol; n=2262; 47 (color) points passing (FCthr=1.5, FdrThrs=0.05
spike
expect at −3.32
− log10 FDR
M−value (log2 fold−change)

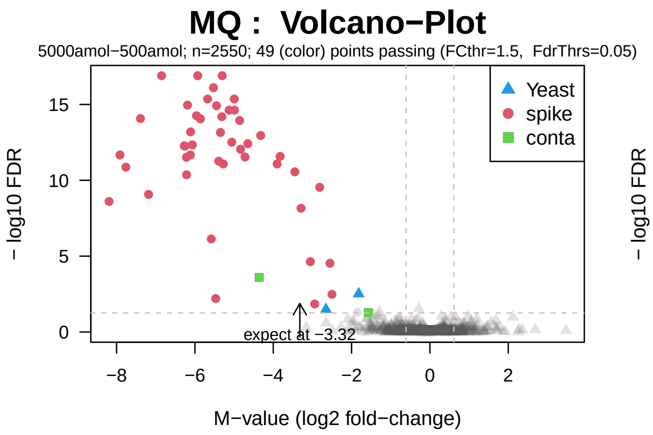
MQ : Volcano−Plot
5000amol−500amol; n=2550; 49 (color) points passing (FCthr=1.5, FdrThrs=0.05)
Yeast
spike
conta
expect at −3.32
− log10 FDR
M−value (log2 fold−change)

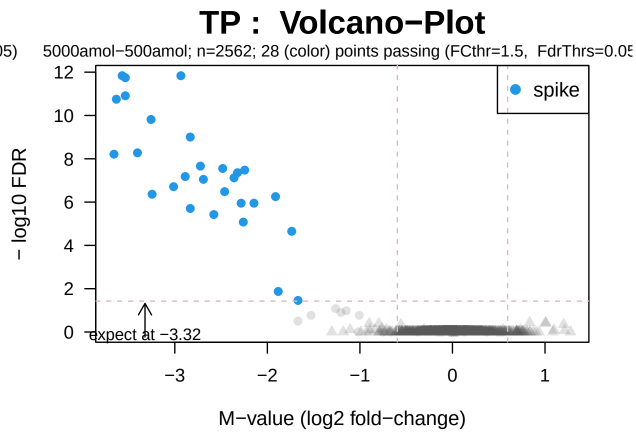
TP : Volcano−Plot
5000amol−500amol; n=2562; 28 (color) points passing (FCthr=1.5, FdrThrs=0.05
spike
expect at −3.32
− log10 FDR
M−value (log2 fold−change)

##### 3.2.3 ROC Curves for the Weakest Group (the '+')

Table 6: AUC details for best pairwise-comparisons

| | Auc.CP | Auc.MQ | Auc.PL | Auc.TP | nCP.BH | nMQ.BH | nPL.BH | nTP.BH |
|---|---|---|---|---|---|---|---|---|
| 500amol-50amol | 0.707 | 0.062 | 0.770 | 0.022 | 22 | 3 | 28 | 0 |
| 250amol-500amol | 0.360 | 0.054 | 0.416 | 0.056 | 3 | 2 | 7 | 0 |
| 10amol-500amol | 0.785 | 0.333 | 0.748 | 0.000 | 23 | 0 | 27 | 0 |
| 10amol-250amol | 0.528 | 0.000 | 0.452 | 0.000 | 14 | 0 | 8 | 0 |
| 10amol-50amol | 0.361 | 0.000 | 0.260 | 0.000 | 5 | 0 | 10 | 0 |
| 250amol-50amol | 0.467 | 0.000 | 0.416 | 0.000 | 12 | 0 | 9 | 0 |
| 100amol-250amol | 0.342 | 0.333 | 0.325 | 0.000 | 6 | 0 | 5 | 0 |
| 10000amol-5000amol | 0.759 | 0.000 | 0.888 | 0.000 | 18 | 0 | 39 | 0 |
| 1000amol-500amol | 0.394 | 0.333 | 0.580 | 0.000 | 9 | 2 | 11 | 0 |
| 100amol-50amol | 0.184 | 0.000 | 0.111 | 0.000 | 3 | 0 | 1 | 0 |

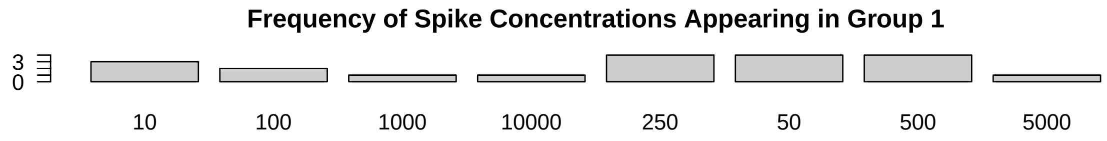


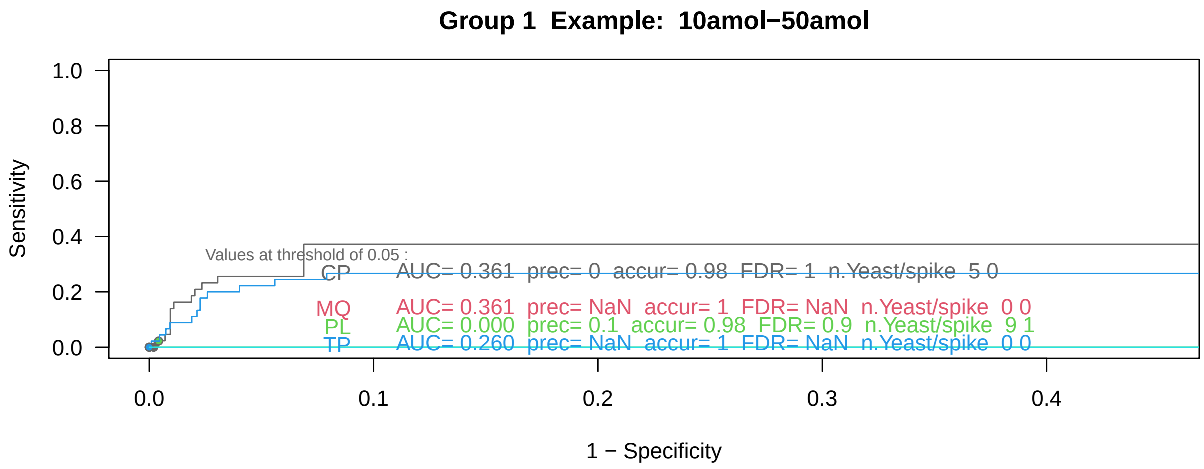

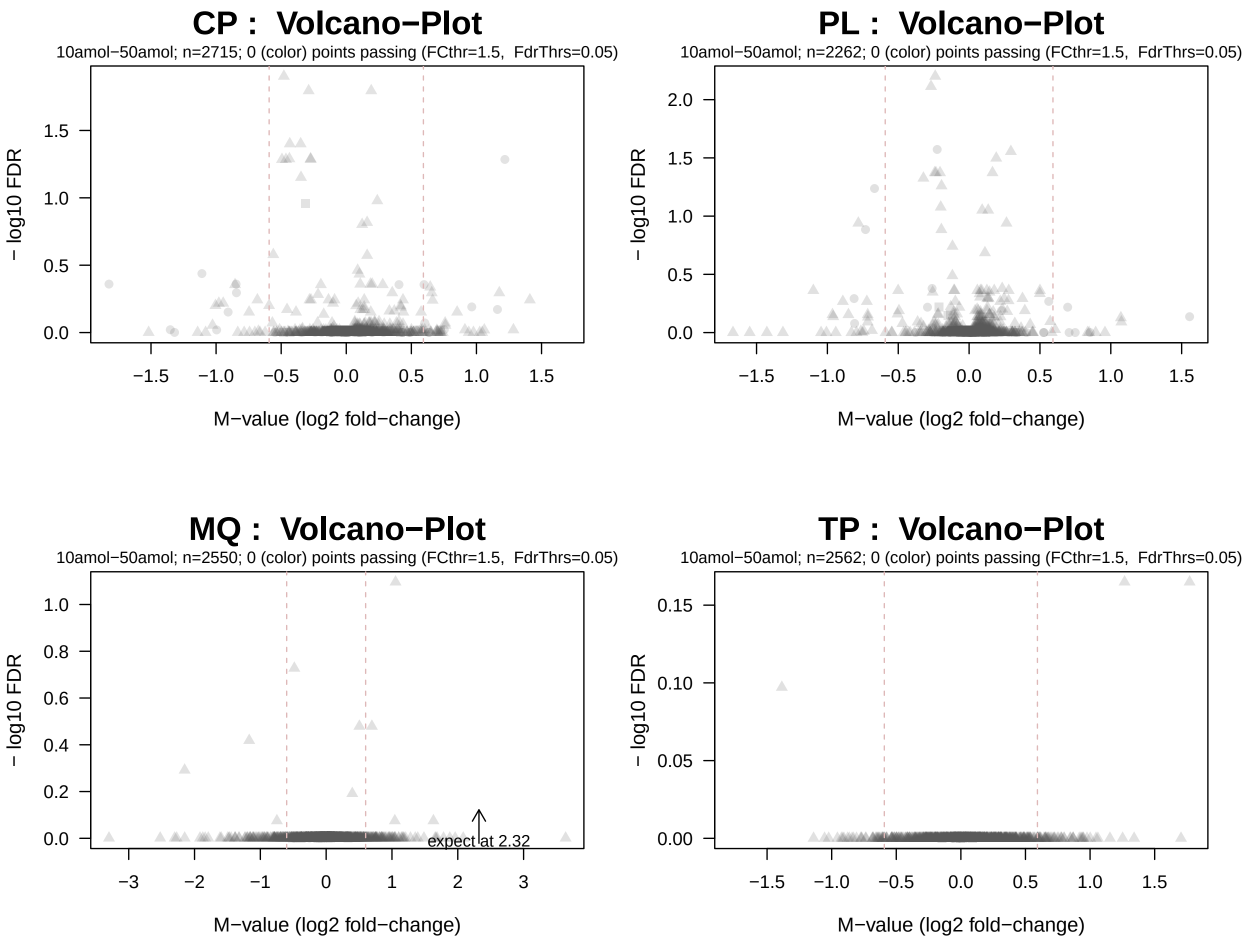
CP : Volcano−Plot
10amol−50amol; n=2715; 0 (color) points passing (FCthr=1.5, FdrThrs=0.05)
PL : Volcano−Plot
10amol−50amol; n=2262; 0 (color) points passing (FCthr=1.5, FdrThrs=0.05)
MQ : Volcano−Plot
10amol−50amol; n=2550; 0 (color) points passing (FCthr=1.5, FdrThrs=0.05)
expect at 2.32
TP : Volcano−Plot
10amol−50amol; n=2562; 0 (color) points passing (FCthr=1.5, FdrThrs=0.05)
− log10 FDR
M−value (log2 fold−change)

### 4 Analysis Focussing On Spike-In Proteins Only (UPS1)

We know from the experimental setup that UPS1 spike-in proteins were added to a constant background of proteins. The samples with the lowest quantities of spike-in added are typically very challenging and it is no surprise that several of the 48 spike proteins were not detected at the lowest concentration(s).

#### 4.1 Global Considerations On All Spike-In Proteins (Number of NAs)

In order to get an overview among the various concentrations of spike-in, let's look how many NAs are in each group of replicates (ie before NA-imputation), and in particular, the number of NAs among the spike-in proteins.

Previsouly we've looked at the total number of NAs, now let's focus just on the spike-in proteins (UPS1). Obviously, instances of non-quantified spike-in proteins make the following comparisons using these samples rather insecure, since NA-imputation is just an 'educated guess'.

In order to have more data available for linear regression modelling it was decided to use spike-in abundance values (UPS1) after NA-Imputation for linear regressions. Previously it was shown that NA values originate predominantly from absent or very low abundance quantitations, which justified replacing NA values by low abundance values in a shrinkage like fashion.

To get started, let's look at the number of NA-values in the various groups of samples.

Table 7: The number of NAs in the spike-in proteins

| | 1 | 2 | 3 | 4 | 5 | 6 | 7 | 8 | 9 | 10 |
|---|---|---|---|---|---|---|---|---|---|---|
| compomics | 50 | 38 | 32 | 16 | 8 | 5 | 2 | 0 | 0 | 0 |
| maxquant | 154 | 175 | 174 | 169 | 133 | 40 | 4 | 3 | 0 | 0 |
| proline | 34 | 17 | 11 | 9 | 0 | 0 | 0 | 0 | 0 | 0 |
| tpp | 180 | 184 | 184 | 179 | 151 | 116 | 44 | 27 | 12 | 3 |

Let's look graphically at the number of NAs in the spike-in proteins along the quantification methods : You can see, some spike-in proteins give more NAs than others.

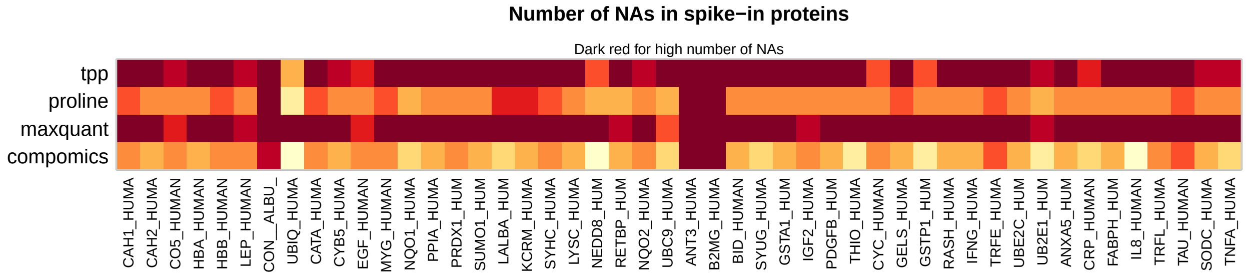


##### 4.1.1 Similarity by PCA (Spike-in proteins only, UPS1)

Plotting the principal components (PCA) typically allows to gain an overview on how samples are related to each other. For more details please see also the vignettes of the R-packages wrProteo and wrGraph. In heterologous spike-in experiments the matrix remains constant, so it makes sense to separate and use ony the spike-in proteins for this type of plot.

As mentioned before, principal component analysis (PCA) cannot handle NA-values. Either all lines with any NAs have to be excluded, or data after NA-imputation have to be used. Here, the option of plotting data after NA-imputation was chosen (in the context of filtering spike-in proteins (UPS1) lines only one whould loose too many lines, ie proteins).

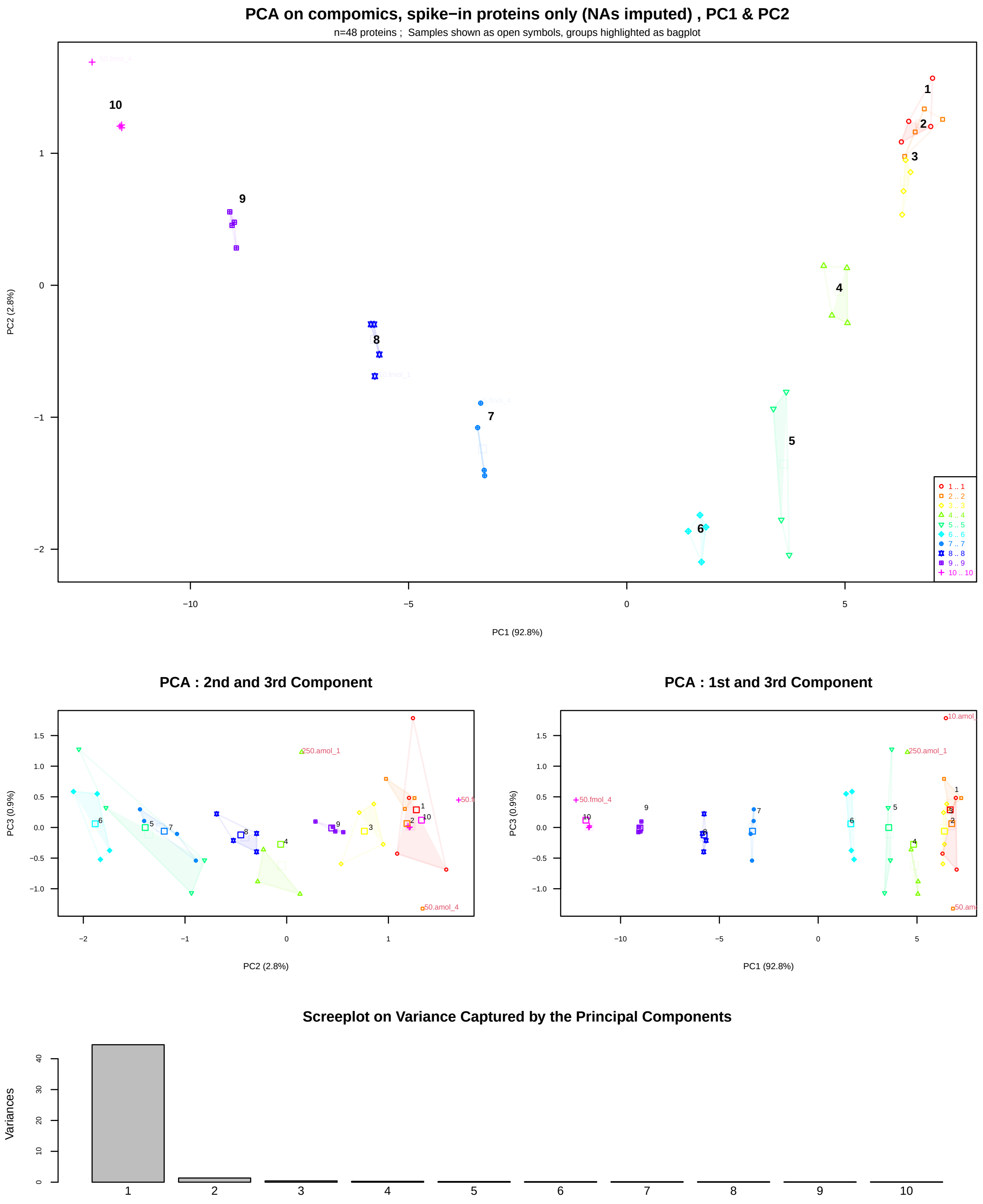

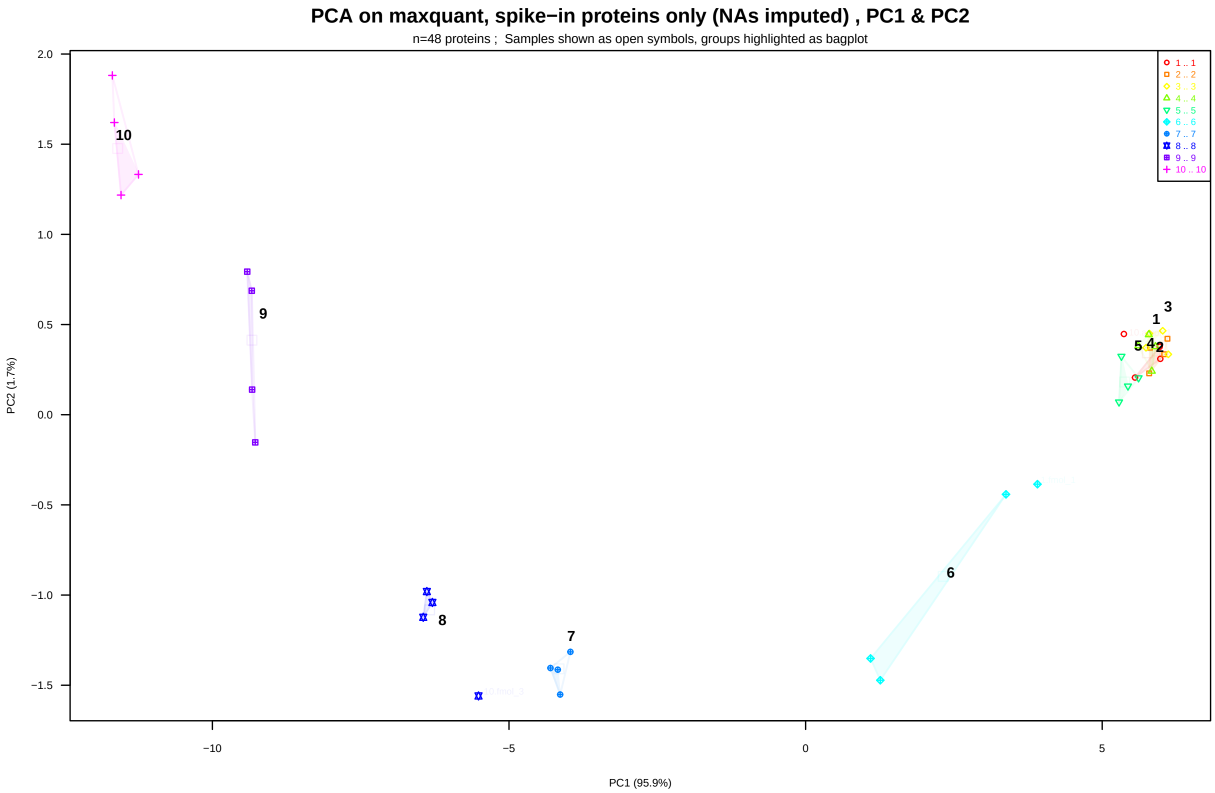
PCA on maxquant, spike−in proteins only (NAs imputed) , PC1 & PC2
n=48 proteins ; Samples shown as open symbols, groups highlighted as bagplot
1 .. 1
2 .. 2
3 .. 3
4 .. 4
5 .. 5
6 .. 6
7 .. 7
8 .. 8
9 .. 9
10 .. 10
PC2 (1.7%)
PC1 (95.9%)

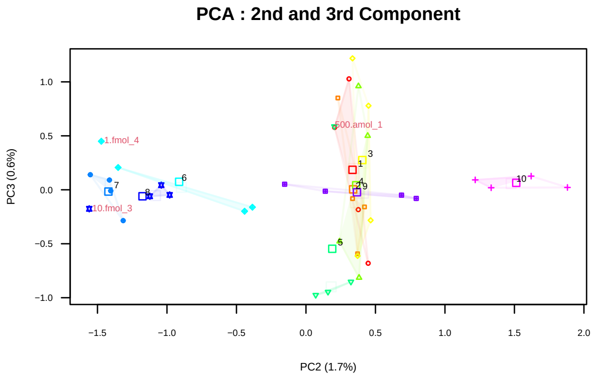
PCA : 2nd and 3rd Component
1.fmol_4
500.amol_1
10.fmol_3
PC3 (0.6%)
PC2 (1.7%)

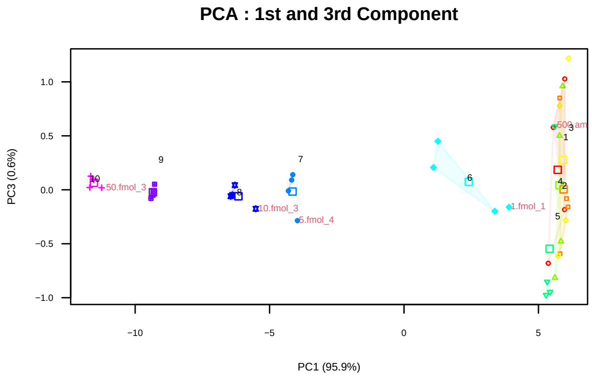
PCA : 1st and 3rd Component
50.fmol_3
10.fmol_3
5.fmol_4
1.fmol_1
PC3 (0.6%)
PC1 (95.9%)

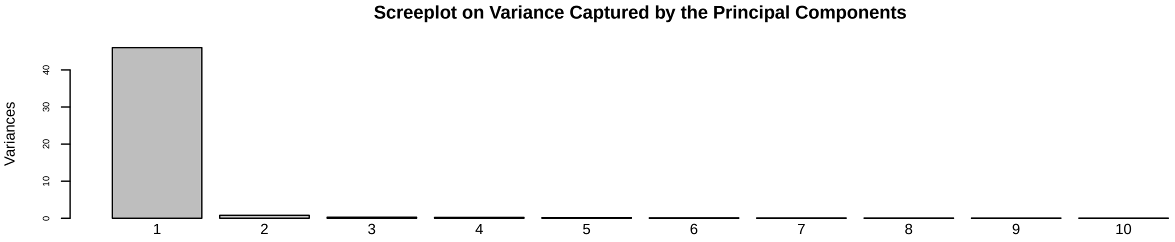
Screeplot on Variance Captured by the Principal Components
Variances

#### PCA on proline, spike−in proteins only (NAs imputed) , PC1 & PC2

n=48 proteins ; Samples shown as open symbols, groups highlighted as bagplot

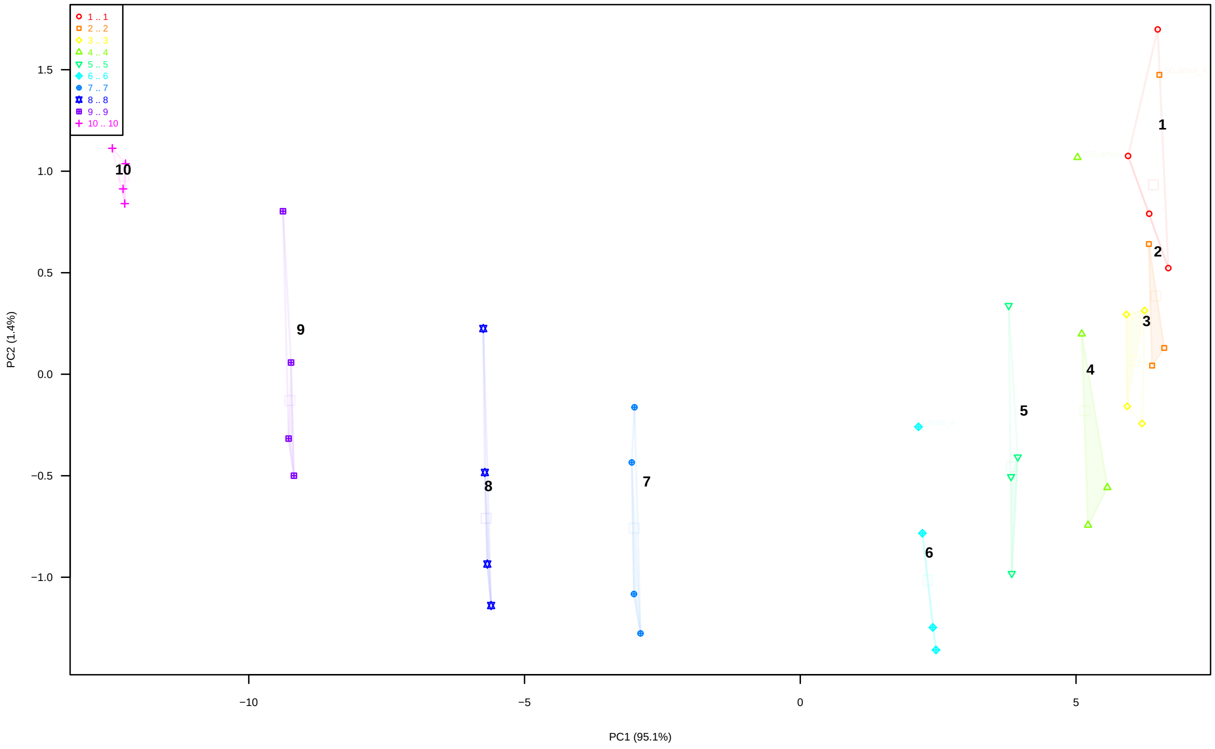


#### PCA : 2nd and 3rd Component

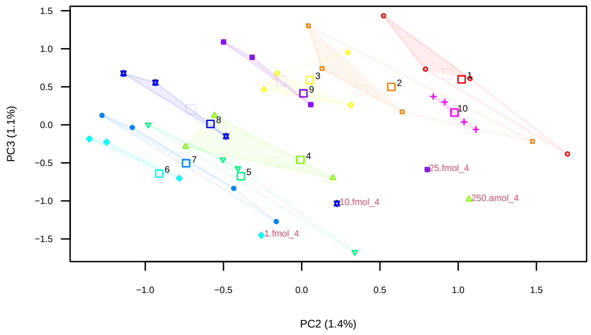


#### PCA : 1st and 3rd Component

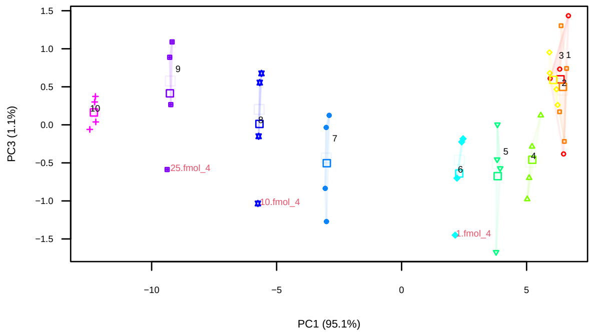


#### Screeplot on Variance Captured by the Principal Components

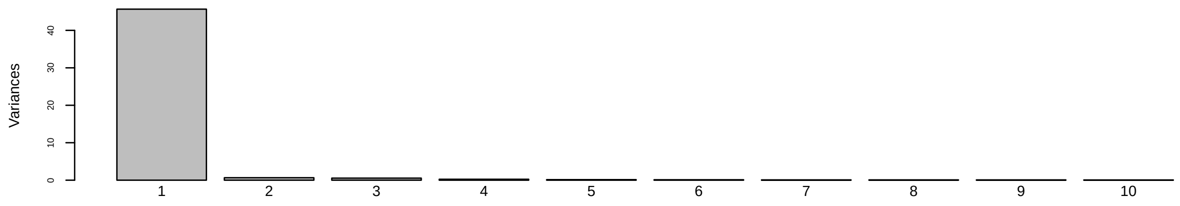

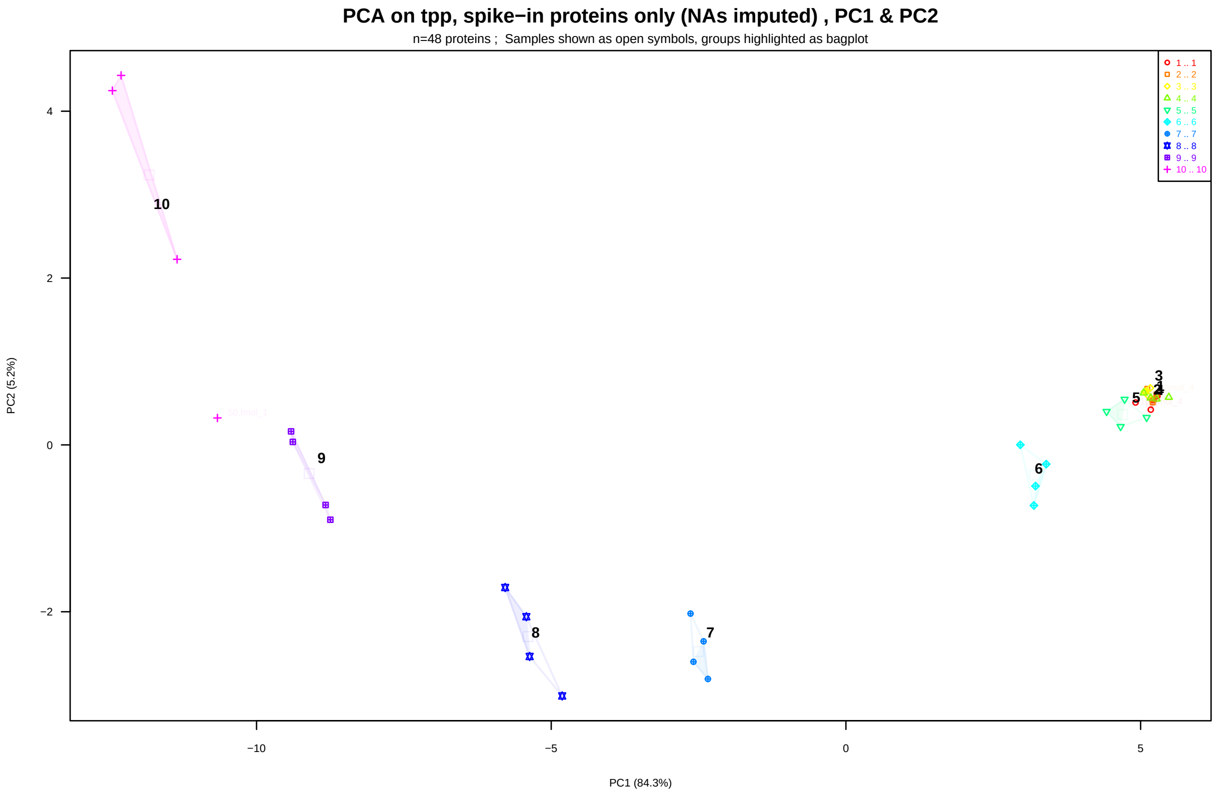
PCA on tpp, spike−in proteins only (NAs imputed) , PC1 & PC2
n=48 proteins ; Samples shown as open symbols, groups highlighted as bagplot
1 .. 1
2 .. 2
3 .. 3
4 .. 4
5 .. 5
6 .. 6
7 .. 7
8 .. 8
9 .. 9
10 .. 10
PC1 (84.3%)
PC2 (5.2%)

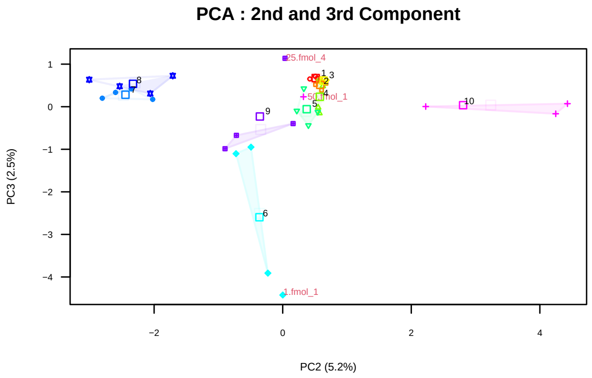
PCA : 2nd and 3rd Component
25.fmol_4
1.fmol_1
PC2 (5.2%)
PC3 (2.5%)

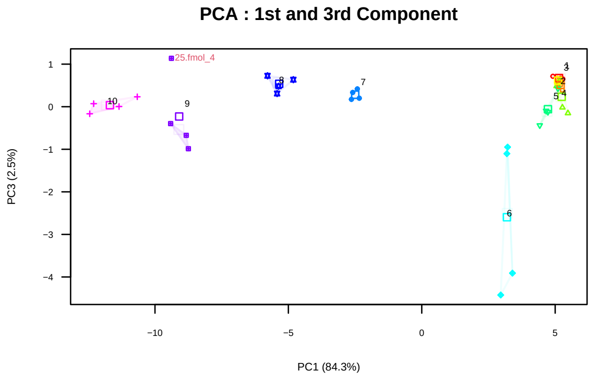
PCA : 1st and 3rd Component
25.fmol_4
PC1 (84.3%)
PC3 (2.5%)

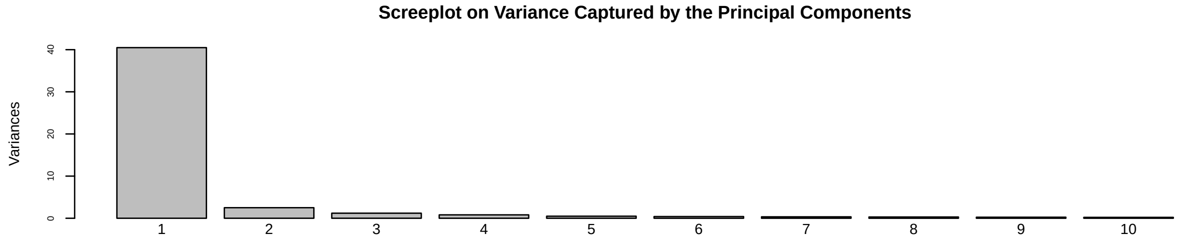
Screeplot on Variance Captured by the Principal Components
Variances

#### 4.2 Testing UPS1 Spike-In Proteins Individually By Linear Regression

In this section we'll look at dose-response curves for all UPS1 spike-in proteins and analyze them by linear regression. Thus, for each spike-in protein we calculate a fresh regression model and extract the slope and p-value (H0: slope=0).

To get an overview, let's look where regressions typically have their best starting-site (ie how many low concentrations points should be better omitted to acchieve best linearity):

Table 8: Frequency of starting levels for regression

| | 10amol | 50amol | 100amol | 250amol | 500amol |
|---|---|---|---|---|---|
| CP | 1 | 7 | 12 | 14 | 14 |
| MQ | 0 | 14 | 13 | 18 | 3 |
| PL | 1 | 6 | 5 | 23 | 13 |
| TP | 1 | 27 | 3 | 16 | 1 |

##### 4.2.1 Summarize Linear Regression Results

When judging results for individual spike-in proteins both the value of the slope as well as the p-value (for H0: slope=0) are important to consider. For example, there are some cases where the quantitations line up well giving a good p-value but with slopes $< 0.4$, although a *slope=1.0* is expected. This is definitely not the type of dose-response characteristics we are looking for.

#### 4.3 Grouping of Spike-In Proteins to Display Representative Proteins

With 48 spike-in proteins plotting all regression lines might not be good choice. Instead, we'll group the regression-results and plot only representative examples for some groups. First we rank all log.p - and abs(1 - slope) -values separately. Thus, for both p-values and the slopes low resulting ranks represent good results and their ranks become 'comparabale'. Then, we (pre)define that we want to organize the data in 5 groups (like restaurant-ratings from 1 to 5 stars), to do so a k-Means clustering approach was chosen.

Previously we organized all spike-in proteins according to their regression characteristics into 5 clusters and each cluster was ordered for descending scores. Now we can use the median position within each cluster as representative example for this cluster.

##### 4.3.1 The Group Of Spike-In Protein Behaving Best (the '+++++')

This group contains the *spike-in* proteins showing the best dose-response curves compared to the expected pattern : These p-values for the regression are nice and the slope is closest to the expected *slope=1* .

Table 9: Regression details for group of the 12 weakest UPS1 proteins

| | Protein | CP slope | MQ slope | PL slope | TP slope | CP logp | MQ logp | PL logp | TP logp |
|---|---|---|---|---|---|---|---|---|---|
| P00915 | CAH1__Carbonic anhydrase 1 | 0.965 | 1.3 | 0.881 | 0.458 | -35.1 | -23.6 | -29.4 | -18.5 |
| P01031 | CO5__Complement C5 | 0.989 | 1.17 | 1.03 | 0.454 | -29.1 | -22 | -36 | -16.8 |
| P04040 | CATA__Catalase | 0.906 | 1.33 | 0.927 | 0.786 | -31.7 | -24.9 | -28.2 | -19 |
| P00167 | CYB5__Cytochrome b5 | 0.958 | 1.3 | 1 | 0.516 | -31.8 | -20.1 | -27.5 | -14.9 |
| P01133 | EGF__Pro-Epidermal growth factor | 0.986 | 0.955 | 0.936 | 0.479 | -27.3 | -21.2 | -29.7 | -16.2 |
| P02753 | RETBP__Retinol-binding protein 4 | 1.12 | 1.12 | 1.11 | 0.642 | -28.9 | -20.2 | -29.7 | -17.3 |
| P16083 | NQO2__Ribosyldihydronicotinamide dehydrog | 0.941 | 1.25 | 0.943 | 0.647 | -29.4 | -19.1 | -34.5 | -18 |
| P01344 | IGF2__Insulin-like growth factor II | 0.98 | 1.13 | 0.976 | 0.464 | -35.7 | -22 | -34.4 | -15.5 |
| P01579 | IFNG__Interferon Gamma | 1.01 | 1.07 | 0.953 | 0.694 | -27.9 | -19.7 | -25.1 | -15.8 |
| P08758 | ANXA5__Annexin A5 | 0.98 | 1.13 | 0.942 | 0.605 | -31.8 | -18.5 | -31.1 | -15.3 |

| | Protein | CP slope | MQ slope | PL slope | TP slope | CP logp | MQ logp | PL logp | TP logp |
|---|---|---|---|---|---|---|---|---|---|
| P10636 | TAU_Microtubule-associated protein tau { | 0.983 | 1.32 | 1.03 | 0.572 | -32.8 | -26.5 | -31.3 | -20 |
| P01375 | TNFA_Tumor necrosis factor, soluble form | 1.05 | 1.24 | 0.954 | 0.301 | -33.9 | -22 | -33.9 | -8.66 |

Now we plot a representative example from the 12 proteins of this group:

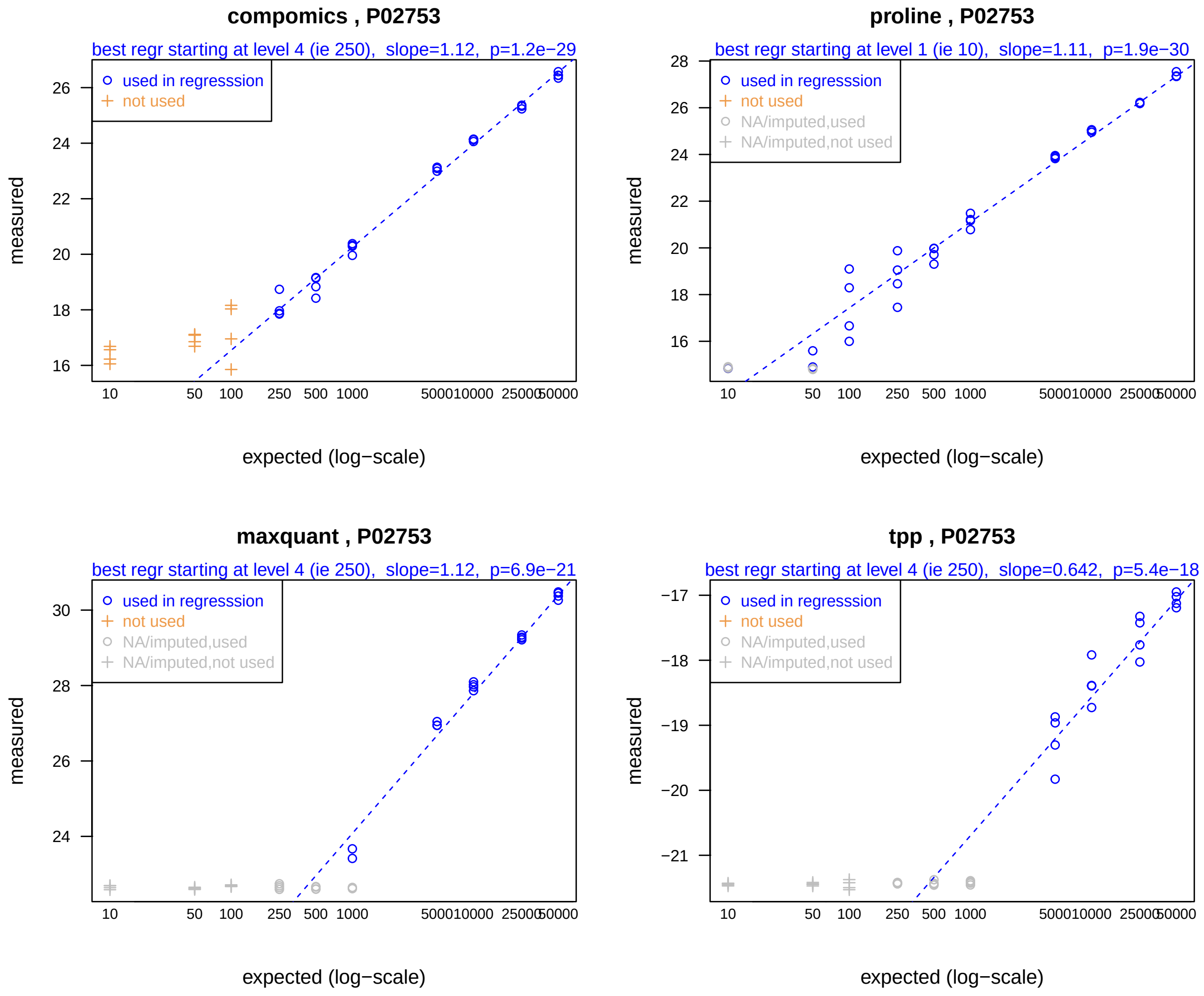


##### 4.3.2 Proteins Of The ‘’Average’‘, ie 3rd Group (the’+++’)

Table 10: Regression details for group of the 13 weakest UPS1 proteins

| | Protein | CP slope | MQ slope | PL slope | TP slope | CP logp | MQ logp | PL logp | TP logp |
|---|---|---|---|---|---|---|---|---|---|
| P69905 | HBA_Hemoglobin subunit alpha | 0.754 | 1.17 | 0.725 | 0.575 | -24.1 | -21.2 | -21.8 | -16.6 |
| P02768 | Albumin | 0.736 | 0.95 | 0.794 | 0.398 | -24.2 | -25.1 | -20.6 | -15.5 |
| P15559 | NQO1_NAD(P)H dehydrogenase | 0.735 | 1.21 | 0.898 | 0.393 | -21 | -20.9 | -25.2 | -8.32 |
| P62937 | PPIA_Peptidyl-prolyl cis-trans isomerase | 0.905 | 1.33 | 0.752 | 0.631 | -28.5 | -17.5 | -25.9 | -17 |

| | Protein | CP slope | MQ slope | PL slope | TP slope | CP logp | MQ logp | PL logp | TP logp |
|---|---|---|---|---|---|---|---|---|---|
| P06732 | KCRM_Creatine kinase M-type | 0.79 | 1.27 | 0.987 | 0.669 | -22.7 | -14.3 | -23.8 | -17.7 |
| P61626 | LYSC_Lysozyme C | 0.817 | 1.03 | 0.792 | 0.285 | -23.6 | -21.7 | -21.6 | -11.3 |
| Q15843 | NEDD8_NEDD8 | 1.15 | 1.19 | 1.03 | 0.147 | -23.7 | -18.6 | -19.1 | -2.34 |
| P63279 | UBC9_SUMO-conjugating enzyme UBC9 | 1.06 | 0.691 | 1.03 | 0.331 | -18.9 | -11.9 | -26.6 | -10.3 |
| O76070 | SYUG_Gamma-synuclein | 0.958 | 1.18 | 0.756 | 0.337 | -24.5 | -19.3 | -22.4 | -10.5 |
| P10599 | THIO_Thioredoxin | 0.914 | 1.35 | 0.918 | 0.282 | -26.8 | -23 | -23.1 | -6.76 |
| P06396 | GELS_Gelsolin | 0.989 | 1.2 | 0.688 | 0.0275 | -29 | -20.3 | -20.9 | -1.93 |
| P10145 | IL8_Interleukin-8, IL-8 | 0.939 | 1.16 | 0.905 | 0.213 | -31.2 | -21.6 | -16.9 | -4.35 |
| P02788 | TRFL_Lactotransferrin | 0.9 | 1.38 | 0.719 | 0.934 | -20.4 | -19.7 | -22.4 | -17.9 |

Now we plot a representative example from the 13 proteins of this group:

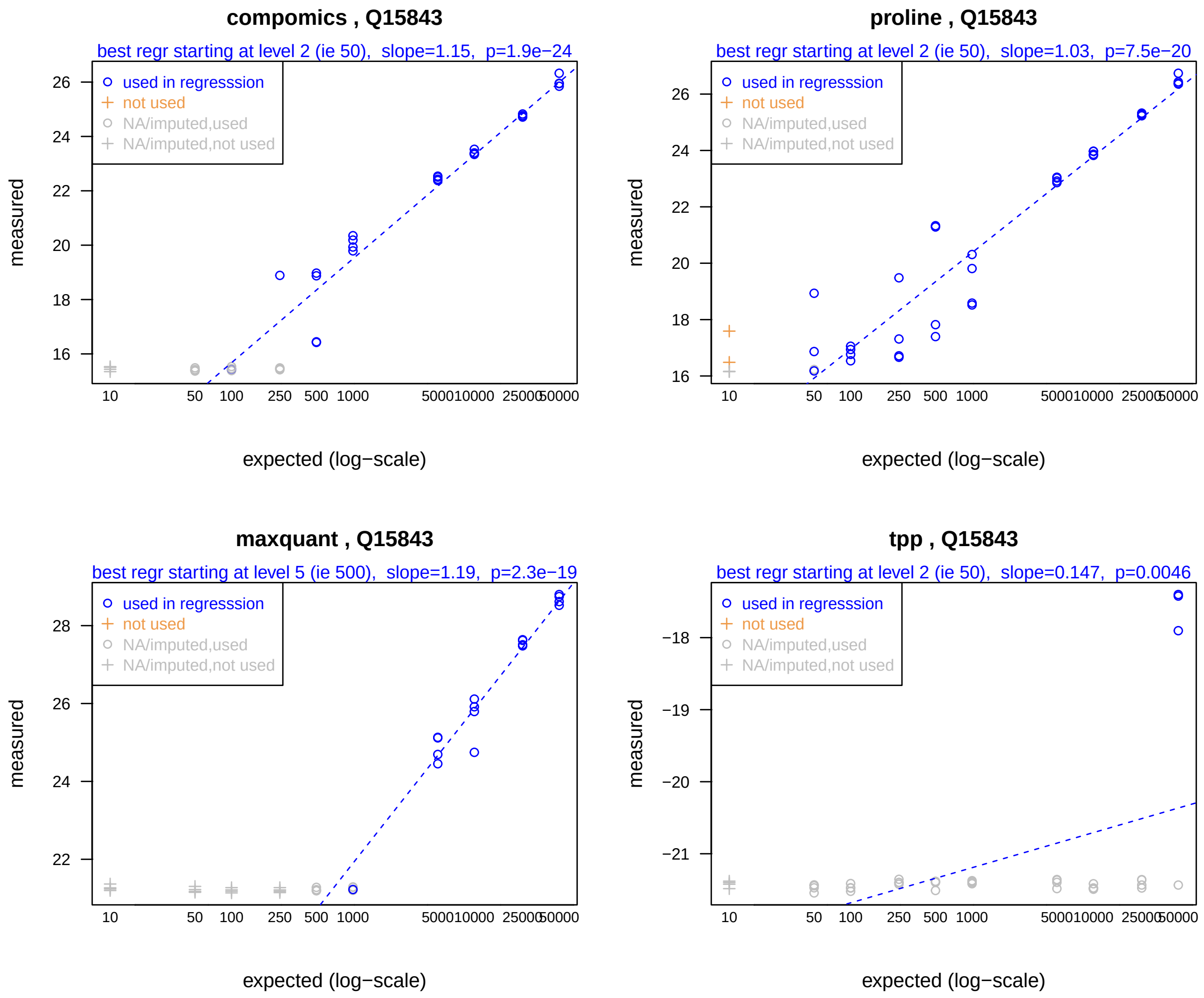


##### 4.3.3 Proteins Of The Weakest Group (the '+')

Table 11: Regression details for group of the 4 weakest UPS1 proteins

| | Protein | CP slope | MQ slope | PL slope | TP slope | CP logp | MQ logp | PL logp | TP logp |
|---|---|---|---|---|---|---|---|---|---|
| P62988 | UBIQ_Ubiquitin | 0.596 | 0.241 | 0.0192 | 0.0374 | -9.62 | -5.74 | -5.27 | -1.46 |
| P61769 | B2MG_Beta-2-microglobulin | 0.52 | 0.654 | 0.551 | 0.291 | -17.2 | -17.6 | -16.1 | -9.05 |
| P08263 | GSTA1_Glutathione S-transferase A1 | 0.755 | 1.37 | 0.311 | 0.325 | -12 | -17.8 | -12.1 | -10.7 |
| P51965 | UB2E1_Ubiquitin-conjugating enzyme E2 E1 | 0.783 | 0.516 | 1.41 | 0.0582 | -9.21 | -11.9 | -22 | -7.36 |

Now we plot a representative example from the 4 proteins of this group:

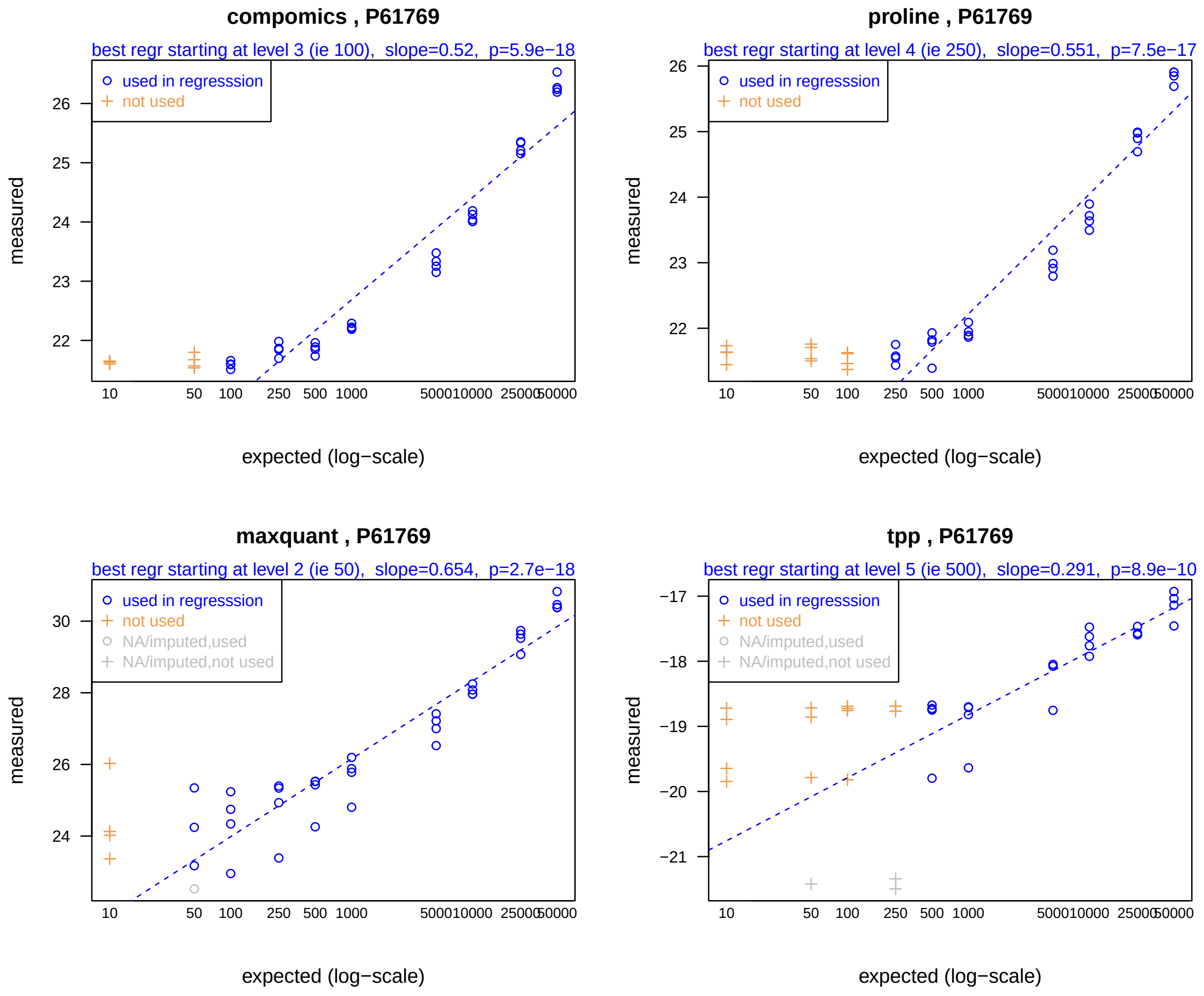


The worst group still contains some dose-response relationships for the higher concentrations of *spike-in* proteins, but many times either the slope differs from the expected *slope=1* or values have high variance.

### 5 Additional Comments

The choice of the 'best suited' approach to quantify and compare proteomics data is not trivial. Most software for quantification of proteomics data do have an important number of 'small' parameters which may have a very strong impact on the final outcome !

Furthermore, ROC curves should also be interpreted with some caution. If a given analysis -software succeeds to identify only 1 out of numerous spike-in proteins, and this protein turns out change abundance 'significantly', the ROC curve will suggest an 'optimal' result. However, typically proteomics users will prefer analysis software finding all spike-in proteins, even if a few false positives from the matrix may be observed.

#### 5.1 Session-Info

For completeness, this document was created using :

```
R version 4.3.1 (2023-06-16 ucrt)
Platform: x86_64-w64-mingw32/x64 (64-bit)
Running under: Windows 10 x64 (build 19045)

Matrix products: default


locale:
[1] LC_COLLATE=French_France.utf8  LC_CTYPE=French_France.utf8
[3] LC_MONETARY=French_France.utf8 LC_NUMERIC=C
[5] LC_TIME=French_France.utf8

time zone: Europe/Paris
tzcode source: internal

attached base packages:
[1] stats     graphics  grDevices utils     datasets  methods   base

other attached packages:
[1] wrProteo_1.10.0.2 wrGraph_1.3.4     wrMisc_1.13.0     knitr_1.43

loaded via a namespace (and not attached):
 [1] gtable_0.3.3       limma_3.56.2       dplyr_1.1.2        compiler_4.3.1
 [5] tinytex_0.46       Rcpp_1.0.11        tidyselect_1.2.0   stringr_1.5.0
 [9] splines_4.3.1      scales_1.2.1       yaml_2.3.7         fastmap_1.1.1
[13] ggplot2_3.4.3      R6_2.5.1           plyr_1.8.8         generics_0.1.3
[17] fdrtool_1.2.17     tibble_3.2.1       munsell_0.5.0      sm_2.2-5.7.1
[21] pillar_1.9.0       RColorBrewer_1.1-3 rlang_1.1.1        utf8_1.2.3
[25] stringi_1.7.12     xfun_0.40          cli_3.6.1          magrittr_2.0.3
[29] digest_0.6.33      grid_4.3.1         lifecycle_1.0.3    vctrs_0.6.3
[33] qvalue_2.32.0      evaluate_0.21      glue_1.6.2         fansi_1.0.4
[37] colorspace_2.1-0   reshape2_1.4.4     rmarkdown_2.24     tools_4.3.1
[41] pkgconfig_2.0.3    htmltools_0.5.6
```
